# Spatiotemporal Atlas of Heart Development Reveals Blood-Flow-Dependent Cellular, Structural, Metabolic, and Spatial Remodeling

**DOI:** 10.64898/2025.12.09.693024

**Authors:** Jooyoung Park, Shuofei Sun, Rohit Agarwal, Andreas Stephanou, Mong Lung Steven Poon, Hyun Maeng, Peyton Lancaster, Iwijn De Vlaminck, Jonathan Butcher

**Affiliations:** Meinig School of Biomedical Engineering, Cornell University, Ithaca, NY, USA

**Keywords:** Hemodynamics, Cardiac morphogenesis, Congenital heart defects, Single-cell transcriptomics, Spatial transcriptomics, Mesoscale, Plasticity, Heart Ventricle, Heart valve, Cell Neighborhood

## Abstract

Embryonic heart development depends on coordinated interactions between cellular programs, molecular signaling, and biomechanical forces, yet how mechanical cues shape cellular and molecular pathways remains incompletely understood. We perturbed cardiac blood flow by partial left or right atrial ligation (LAL/RAL) in chick embryos, generating chamber-specific hemodynamic gain- or loss-of-function states. Using single-cell and unbiased, high-resolution spatial transcriptomics, we generated a spatiotemporal atlas of flow-dependent tissue development, enabling systematic investigations of bidirectional interactions between blood-flow mechanics and tissue development. Spatially resolved analyses recapitulated key features of normal morphogenesis including regional maturation and the cellular neighborhood. We further revealed flow-specific remodeling across molecular, cellular, and architectural levels. Altered flow induced LOX-expressing cardiomyocyte and endocardial states, disrupted ventricular layer organization, and delayed maturation, alongside transient metabolic and ion-transport adaptations. Together, these findings define how redistributed blood flow reshapes developing cardiac tissues and provide a framework for studying flow-dependent remodeling in morphogenesis and malformation.

## INTRODUCTION

The heart is the first functional organ to form in the embryo and one of the organs most profoundly shaped by biomechanical forces, particularly hemodynamic load, which both influence and are influenced by tissue growth^1^. Disruption of this bidirectional coupling alters ventricular lumen dimensions and the balance between trabecular and compact myocardium, contributing to severe congenital heart defects such as hypoplastic left heart syndrome (HLHS) and left ventricular non-compaction (LVNC). Although extensive genetic studies have sought to disentangle the complex continuum of developmental cardiac phenotypes, genotype-phenotype correlations remain limited, underscoring that genetic programs alone cannot fully account for the spectrum of developmental cardiac phenotypes. These limitations highlight the equally critical role of non-genetic factors, especially biomechanical cues, and the intimate interplay between mechanical forces and cellular and molecular signaling during heart morphogenesis^2–6^, a relationship that remains incompletely understood. Motivated by this, we aimed to define how mechanical forces regulate cellular patterning and tissue remodeling during heart development using an integrated framework that combines controlled redistribution of blood flow with single-cell and spatial transcriptomics.

Here, we performed LAL and RAL in growing embryos to redistribute blood flow and asked how altered hemodynamics shapes cardiac development at the molecular, cellular and local tissue-architectural levels. Using a small eggshell window, we conducted microsurgery and resealed the shell to allow continued growth. Prior studies with this approach typically maintained survival to embryonic day 7 due to the drop of viability with the completion of ventricular and OFT septation. This challenge limits analysis to early stages when many structures remain immature^7–9^. By finding a location of the surgical loop not obstructing the rotations of the outflow tract, we increased viability, extending survival to day 12. This innovation enabled us to analyze at later stages with more defined cardiac morphology.

From pre- and post-surgical time points, we collected hearts under distinct hemodynamic conditions, including sham control (hereafter referred to as normal), LAL, and RAL, and built a spatiotemporal and hemodynamically regulated cardiac cellular and molecular atlas by integrating single-cell RNA sequencing with unbiased, high-resolution spatial transcriptomics. Specifically, we mapped cellular phenotypes, cell plasticity which implicates cell state, examined spatial layer ontology, and elucidated emergent cellular phenotypes in abnormally remodeling ventricles. Together, these results outline the mechanobiological–molecular interplay in cardiac morphogenesis and provide a framework to identify signatures and regulatory pathways relevant to congenital malformations.

## RESULTS

### A Spatiotemporal Single-cell and Spatial transcriptomics Atlas of the Developing Fetal Chicken Heart Under Different Hemodynamics

To test how blood flow affects heart tissue development, we reduced the atrioventricular preload of the left or right ventricle via partial-LAL or RAL, which simultaneously increases the preload to the contralateral ventricle via blood shunting through the atrial foramen ovale (**Figure 1A**). We performed this surgery after initial cardiogenesis (HH24/Day 4) and maintained the hemodynamic condition until HH40/Day 12, after the major morphogenic events of heart development were complete. This experimental design allowed us to test the cellular, molecular, and morphogenetic consequences of chamber-specific hemodynamic gain- and loss-of-function.

**Figure 1.**
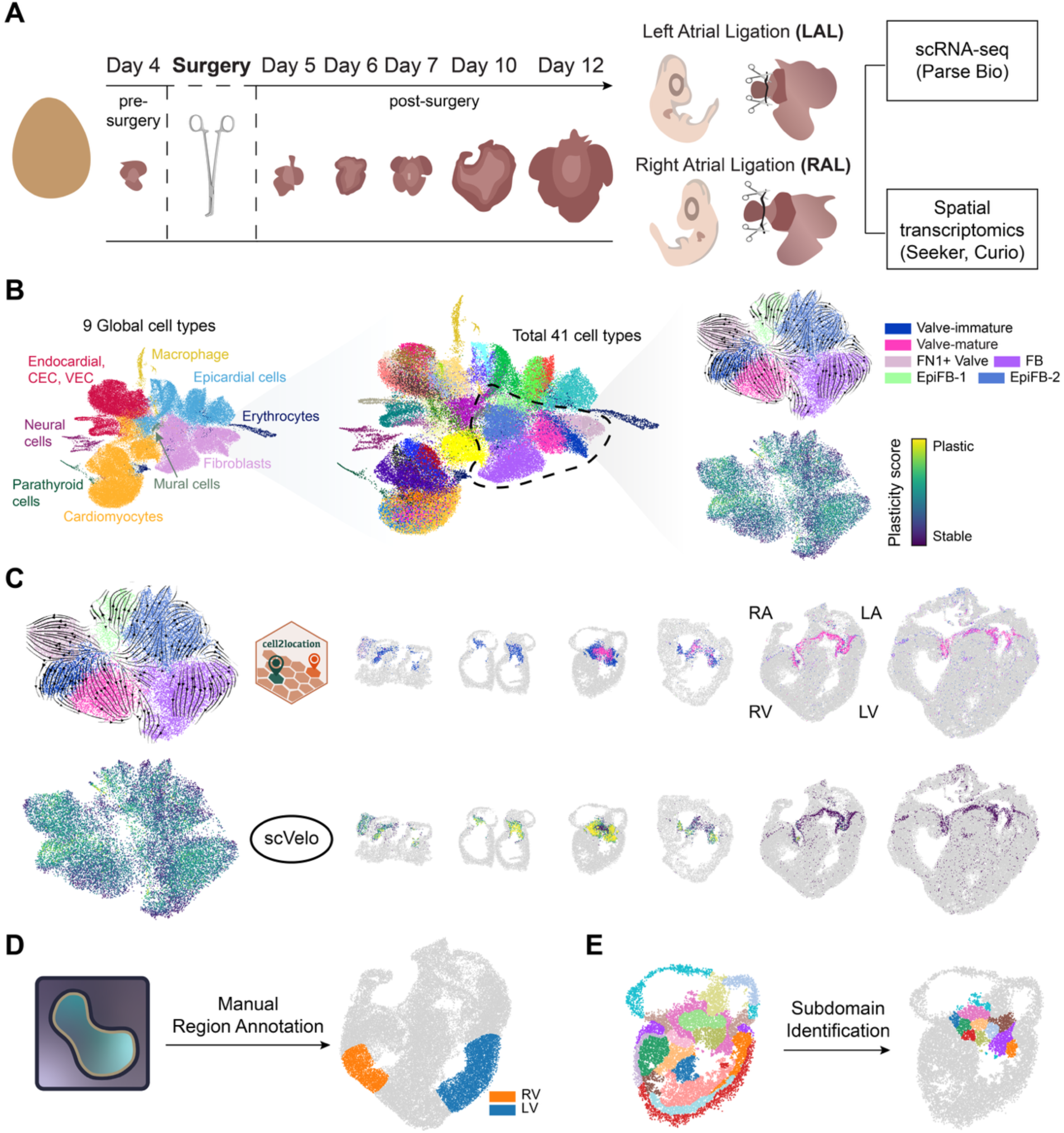
Spatiotemporal single-cell and spatial transcriptomics atlas of the heart under perturbed hemodynamics. (A) Schematic overview of sample collection and surgical perturbation. Heart samples were collected at multiple developmental stages and processed for single-cell or spatial transcriptomic profiling. Hemodynamic perturbations were introduced by left atrial ligation (LAL) or right atrial ligation (RAL). (B) Workflow of scRNA-seq analysis. Cell type annotation was performed hierarchically, beginning with coarse classification followed by granular subtypes. RNA velocity analysis was carried out independently within each coarse cell type. (C) Overview of cell type and plasticity score mapping. Cell types were assigned to spatial spots using cell2location based on the dominant contributing cell type. Plasticity scores were computed for each spot as the product of the cell type proportion and its corresponding cell type plasticity score. (D) Schematic of manual region annotation with Napari for left ventricle (LV) and right ventricle (RV) analysis. (E) Schematic of subdomain identification for valve neighborhood analysis with recursive GraphST.

We combined single-cell transcriptomics (Split-seq, Parse Biosciences) and high-resolution spatial transcriptomics (Seeker, Curio) across fetal chick cardiac morphogenesis from post-heart formation (HH24/Day4) to late fetal morphogenesis (HH40/Day12) (**Figure 1A**), including key milestones: completion of ventricular septation (HH30/Day7), and coronary artery network formation (HH36/Day 10). Single-cell RNA sequencing (scRNA-seq) was performed on normal hearts (sham surgical controls) and hearts for which ventricles were hemodynamically de-loaded or over-loaded (LAL and RAL surgeries) at day 4 and followed to day 5 (24 hours post-surgery) and subsequent stages. We integrated single-cell transcriptomes across all hemodynamic conditions and developmental stages, yielding 80,197 cells and enabling construction of a detailed temporal single-cell atlas (scAtlas) of the developing biventricular heart under hemodynamic perturbation. We initially grouped cells into nine major lineages, including epithelial cells, fibroblasts, endothelial cells, and cardiomyocytes, followed by fine-grained subclustering to resolve cellular subtypes. This analysis identified both canonical cardiac populations and subtypes enriched in perturbed hearts (**Figure 1B, S1, S4A, B; STAR Methods**). For spatial transcriptomics (ST), we collected data from normal hearts at days 4, 5, 6, 7, 10, and 12; from LAL hearts at days 5, 6, 7, 10, and 12; and from RAL hearts at day 10 and 12 (**Figure 1A; STAR Methods**). Each sample contained between 8,467 and 67,681 spots, with the number of spots increasing at later developmental stages.

Leveraging the matched scRNA-seq and high-resolution ST datasets, we constructed an integrated cell-type and cell-state plasticity map across normal and perturbed cardiac development (**Figure 1C**), characterize changes in ventricular wall architecture and molecular composition under altered hemodynamic load (**Figure 1D**), and delineate spatially distinct subdomains within the developing heart valves (**Figure 1E**). Together, this approach enabled us to resolve dynamic cellular transitions over time, revealing the cellular neighborhoods and interactions that coordinate chamber maturation and valve morphogenesis.

### High-Resolution Spatial Map Reveals Normal Heart Development Processes

We first investigated normal fetal heart development. We analyzed scRNA-seq data to annotate cell types and perform RNA velocity analysis which characterized cell-state transitions in the developing hearts and was used to derive a plasticity score for each cell type (**Figure S1; STAR Methods**)^10^. This plasticity score reflects a cell’s potential for state transition, with higher scores indicating cells in a more dynamic, transitional state, and lower scores reflecting cells in a more stable state. Using cell2location, we mapped cell type compositions onto ST spots, assigning each spot to its most dominant cell type^11^. We then computed a spatial plasticity score as the proportion-weighted sum of cell-type plasticity scores per spot (**Figure 1C; STAR Methods**). This enabled the spatiotemporal mapping of developmental cell states and structural organization within specific tissue populations.

### Cellular evolution during fetal atrioventricular valve morphogenesis

The prevalvular atrioventricular (AV) cushions maintain unidirectional blood flow within the developing heart as they fuse, extend, and condense into valve leaflets^12^. Our spatial atlas revealed that the early day 4 AV cushion is predominantly composed of Valve-immature (TBX20+/SOX5+) mesenchymal cells. The outflow cushion in contrast is predominantly composed of FN1+ Valve mesenchyme (FN1+/ROBO1+) (**Figure 2A**). Valve-immature and FN1+ valve cells persist in the day5 AV cushion. Interestingly, after AV cushion fusion (HH27/Day 6), Valve-mature fibroblasts (COL1A2+/COL6A2+) emerge in the interior of the AV cushion mass while the leaflet tips remain predominantly Valve-immature. The Valve-mature phenotype progressively dominates the valve, with near-complete transition observed by days 10 and 12. This maturation process is accompanied by a decline in spatial plasticity, with elevated plasticity scores observed around days 5-6 in valve regions undergoing active cell state transitions (**Figure 2A**). We further identified an evolution in atrioventricular valve endothelial phenotype (VEC). Initially, TBX20+ VEC-immature (ITPR2+/FGFR3+) were detected, while VWF⁺ VEC (COL25A1+/PLCB4+) were very scarce (**Figure 2G**). From day 7 onward, TBX20+ VEC-immature cells dwindled in proportion but remained localized to the valve periphery in close proximity to Valve-immature mesenchyme. Mature VWF+ VEC (COL25A1+/PLCB4+) then prominently established the distinct endothelial boundary surrounding the left and right AV valve structures by day 10 (**Figure 2A, E**). Interestingly, we observed a residual cushion mass on the right atrioventricular valve (RAV) apparatus at day 10, which contained a mixed population of immature and mature valvular interstitial cells (VIC) enclosed by TBX20⁺ VEC-immature cells (**Figure 2G)**. Notably, this local area exhibited a high plasticity score despite the rest of the valve being highly matured (**Figure 2A**). Taken together, these findings identify a progressive maturation of the AV valve phenotype concomitant with their structural morphogenesis, with the morphing elements exhibiting higher levels of plasticity. The temporal wave of maturation progresses from the base to free edge of the leaflets, converting both VIC and VEC phenotypes.

**Figure 2.**
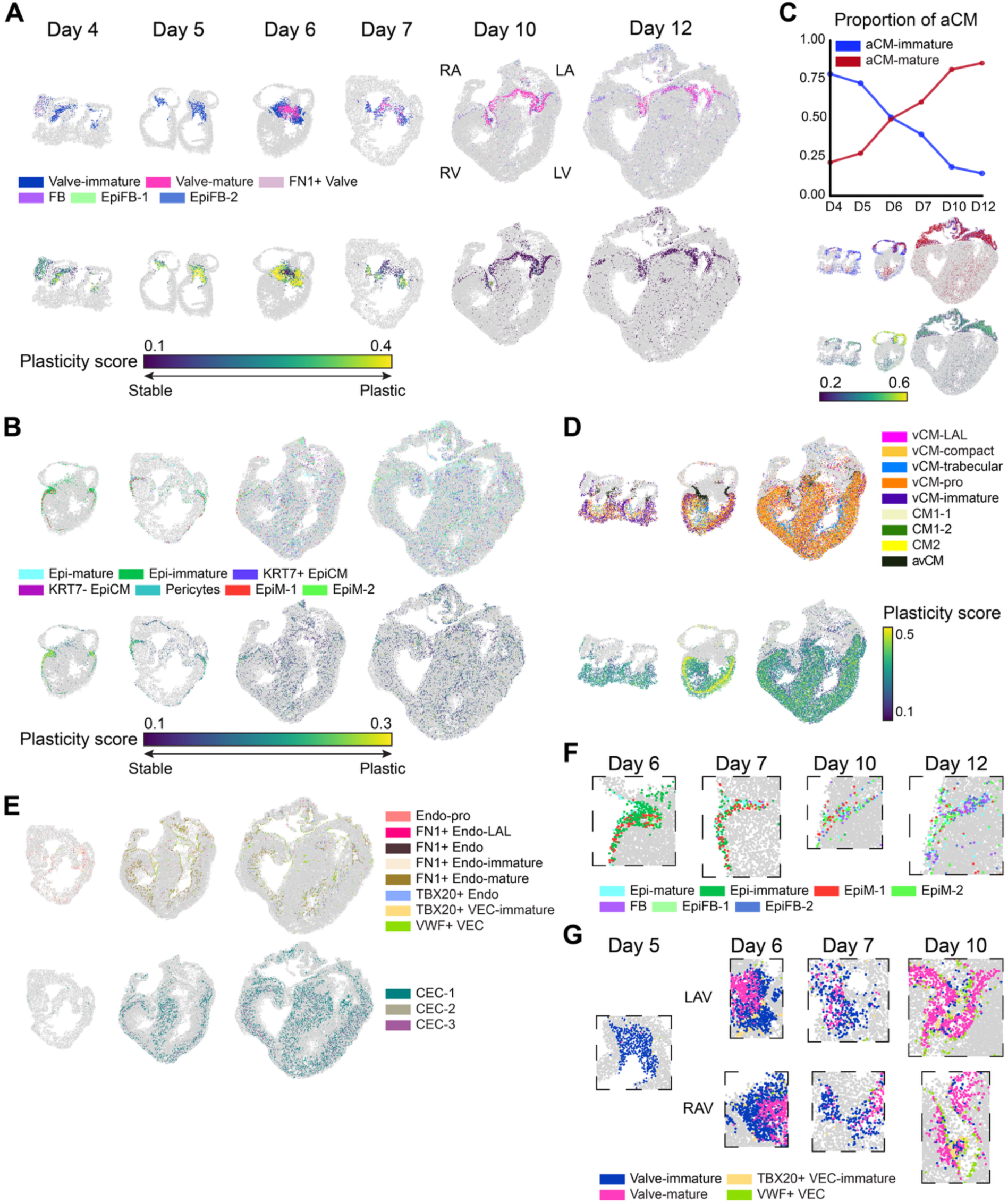
Cellular dynamics during normal heart development. (A) Cell type map (top) and plasticity score map (bottom) of fibroblasts across developmental time points (Day 4, 5, 6, 7 and 12). (B) Cell type map (top) and plasticity score map (bottom) of epithelial cells across five time points (Day 6, 7,10 and 12). (C) Line plot of proportional changes in aCM-immature and aCM-mature across developmental time points (Day 4, 5, 6, 7 and 12). Representative maps of cell type and plasticity score shown for Day 4, Day 6, and Day 12. (D) Cell type map (top) and plasticity score map (bottom) of vCMs, CMs, and avCM across timepoint Day 4, 6, and 10. (E) Cell type map of endothelial cells at Day 7 and 10, displaying endocardial cells and VECs (top), CECs (bottom). (F) Zoomed in a spatial map around the ventricular shoulder. (E) Zoomed in a spatial map around the valve. LAV: left atrioventricular valve, RAV: right atrioventricular valve

### Epicardial derivation of the ventricular shoulders and interventricular mesenchyme

The primitive epicardium first populates the exterior of the heart via migration from the pro-epicardial organ^13,14^. We identified Epi-immature cells as forming this at day4/5. Epicardial-to-mesenchymal transition (EMT) then converts epicardial cells into mesenchymal-like progenitors contributing multiple non-cardiomyocyte lineage populations^15^. We first observed evidence of EMT at day 6, marked by the emergence and expansion of mesenchymal-like epicardial cells EpiM-1 (NPAS3+/LTBP1+) and EpiM-2 (KCNQ5+/FREM1+) (**Figure 2B, F, S2**). At day 7, KRT7+ EpiCM (IQSEC1+/SNTG2+) further emerge at the shoulders of the ventricles, in close proximity to EpiM-1 and EpiM-2 and spatially coincide with regions exhibiting elevated plasticity scores, consistent with active cellular transitions. Epi-immature cells begin to dissipate at this time in favor of Epi-mature (IGFBP5+/PTPRF+), with mesenchymal EpiM populations becoming dominant in these shoulder regions. Adjacent to these cells, we further identified the emergence of epicardial-like fibroblasts EpiFB-1 (SYNE2+/BNC2+) and EpiFB-2 (SLIT3+/DLG2+) at day 10. Non-valve fibroblasts have previously been reported to originate from the epicardium. These populations likely represent epicardial-derived fibroblasts^16^. By day 12, FB (FBLN5+/LTBP2+) fibroblasts (**Figure 2B, F, S2; Table S1**) appear. This region undergoing EMT, together with the emergence of multiple mesenchymal cell types, is consistent with the progressive formation of the fibrous sheath that separates the conduction of the atrium and ventricle. The epicardium further contributes non-myocyte mesenchyme in the ventricular interior^13^. Non-myocardial mesenchymal cells were not detected until day 6, with some pericytes (POSTN+/EBF1+) emerging within the ventricular myocardium (**Figure 2B**). Mesenchymal populations remained sparse until day 10, when multiple epicardial derived cells emerged throughout the ventricles, including EpiM-1, EpiM-2, and KRT7+ EpiCM. Pericytes are known to support ventricular homeostasis through interactions with endothelial cells and cardiomyocytes, and play an essential role in coronary vessel maturation, as the capillary endothelial plexus cannot properly resolve in their absence^17^. The spatial distribution of pericytes in our map supports the establishment of robust coronary vascular networks during later stages of development (**Figure 2B, S2**). These results establish a progressive population of the cardiac ventricles by epicardial derived mesenchymal progenitors, who subsequently differentiate into fibroblasts and pericytes to support atrioventricular conduction insulation and coronary vascular maturation.

### Atrial Myocardial Maturation Dynamics

The right and left atrial chambers are composed of a distinct population of cardiomyocytes known as atrial cardiomyocytes (aCMs). The contraction of the early heart tube is facilitated by atrial-like myosins, which are replaced with ventricular myosins with the formation of the ventricular segments. While the precise timing of atrial myocardium maturation is unclear, previous studies suggested it begins around day 5^12^. Consistent with this, we detect atrial cardiomyocytes in the early atrial and ventricular segments at day 4 (**Figure 2C, S2**). At this timepoint, the early embryonic heart has a single atrial segment, in which we identified aCMs as dominantly aCM-immature (PTN+/TTN+), interestingly with limited plasticity. These cells then differentiate to a more mature aCM-mature (MYH7+/TTN+) phenotype starting at day 5. In contrast to the atrioventricular valves, this transition initially increases in atrial plasticity from day 6, indicating active differentiation dynamics (**Figure 2C**). Interestingly, this maturation appears to initiate in the region of the atrial septum, while the free walls of the atria remain aCM-immature. Further, aCM-mature cells concentrate within the early ventricular septal protrusion, while the residual aCM-immature cells of the ventricles remain in the free wall. Atrial maturation continues from the center outward from day 6 to day 12, until virtually all of the atria are aCM-mature phenotype. At the same time, the density of atrial CM in the ventricles drops precipitously and the remaining cells convert to the aCM-mature phenotype by day 12. No cellular patterning differences were noted between the right and left atrium. These results establish a more specific spatial and temporal evolution of the maturation process, with day 6 being a prime time for active transition. This period is concomitant with the formation of the atrial septum, separation of left and right ventricular inflows, and establishment of distinct left and right atrial chambers.

### Spatial Ventricular Differentiation Dynamics

Beneath the atria lie the two lower chambers of the heart, the ventricles. The early embryonic heart has a single ventricular segment. We identify ventricular cardiomyocytes (vCMs) first appear as early-stage vCM-immature (UNC5C+/DACH1+) at day 4, predominantly located in the outer layer (**Figure 2D, S2**). These cells are likely synonymous with “primary” ventricular cardiomyocytes^18^. As development progresses, more mature subtypes emerge. vCM-compact (RYR+/MYH15+) and proliferative vCM-pro (CENPF+/TOP2A+) localize to the putative compact layer of the ventricle, while vCM-trabecular (PDE1C+/CORIN+) are found in the inner layer of the chamber and the initial ventricular septal protrusion. Interestingly, the plasticity score indicates that the trabecular region of the ventricular chambers is more stable than the compact domain (**Figure 2D, S1, S2**). This is consistent with prior reports localizing proliferation in the compact myocardium from day 8 to day 14, after completion of ventricular septation^19^. Additionally, we identify atrioventricular junctional myocardial cells avCM (HS6ST3+/PDE1C+) adjacent to the AV valve region (**Figure 2D**). These cells are prominent during AV cushion formation and fusion and remain present at the myocardial junction to which the AV valvular apparatus attaches. The single cell data further identifies three additional CM subtypes, which we denote CM1-1, CM1-2, and CM2 based on their transcriptional profiles, where they are generally distributed in the ventricle of the hearts (**Figure S1, S2 and Table S1**). Further, these populations compositions and spatial localization are largely consistent between left and right ventricles, supporting their morphogenesis is similar at the cellular level even though the ventricular walls acquire different volumes of each compartment largely consistent with their hemodynamics.

### Spatial Evolution of the Endocardium and Coronary Vascular Network

The early embryonic heart has no internal vascular supply. The initial coronary capillary plexus is formed around day 8 of chick heart development via ingress from sinus venosus and endocardial derived progenitors^20^. These cells dedifferentiate and differentiate into arterial and venous phenotypes^21^. By day 12, the coronary arterial and venous trees are fully established^12^. Consistent with these morphological milestones, our data identifies coronary endothelial cells (CECs) emerging predominantly at later stages, beginning at day 6 (**Figure 2E, S2**). We identified three CEC subtypes. CEC-1 (FLT1+/PLXND1+) appears first and appears to migrate caudally into the ventricular space. CEC-2 (RELN+/NFATC1+) and CEC-3 (MYH15+/RYR2+) appeared later (day 10), with all three CECs broadly distributed in the ventricular walls by day 12, similar to the expansion of epicardially derived mesenchymal cells. These results suggest that CEC-1 and CEC-2 are consistent with these coronary artery origins and further support the timing of the expansion and maturation of the coronary tree.

In contrast, endocardial cells were detected across all time points, although their subtype composition and spatial organization evolved over time (**Figure S2**). Endocardial progenitors from endoderm co-form the inner lining of the initial heart tube while the primary cardiomyocytes form an outer contractile sleeve^22^. During the early fetal stages (from day 4 to 7), proliferative Endo-pro (TOP2A+/CENPF+) and FN1⁺ Endo-immature (FN1+/SNTG2+) were most abundant and exhibited no clear spatial confinement (**Figure 2E, S2 and Table S1**). These early endocardial progenitors ECM components such as FN1, which support the formation of the endocardial lining^23,24^. At later stages, the spatial organization of endocardial populations became more defined, forming a distinct inner layer of the ventricular wall and lining the trabecular protrusions (**Figure 2E, S2**). These lining cells include FN1+ Endo-mature (FN1+/COL4A5+) and TBX20+ Endo (FGFR3+/ITPR2+). FN1+ Endo-mature appears to be more prominent at day 10 while TBX20+ Endo proportionally more at day 12, particularly at the base of the trabeculae (**Figure S2**). A FN1+ Endo population was also identified in the single cell data but was not localized in the spatial data (**Figure S1, S2 and Table S1**). We did not find significant differences in spatial or temporal endocardial patterning with respect to left and right ventricles. These results significantly augment our understanding of the spatial and temporal evolution of endocardial development in the heart.

Collectively, our high-resolution spatial cell type atlas not only recapitulates the key cellular and structural processes underlying normal heart development but also delineates the specific cell types contributing to these events. This underscores the utility of spatial transcriptomics in resolving the complex heterogeneous cellular dynamics that shape cardiac morphogenesis.

### Emergence of LOX Expressing LAL-enriched Cell types in Right Ventricle of LAL Heart

Using scRNA-seq analysis, we identified two cell types, ventricular cardiomyocytes (vCM-LAL) and endocardial cells (FN1+ Endo-LAL), that are enriched in LAL hearts and largely absent in other conditions (**Figure 3A, B, S4A, B**). Among differentially expressed genes (DEG) for these LAL-specific populations, we found *LOX* (Lysyl oxidase) as a shared marker gene (**Table S1 and Figure S4A, B**). These findings prompted spatial analysis of their distribution in LAL hearts and assessment of whether *LOX* expression distinguishes conditions.

**Figure 3.**
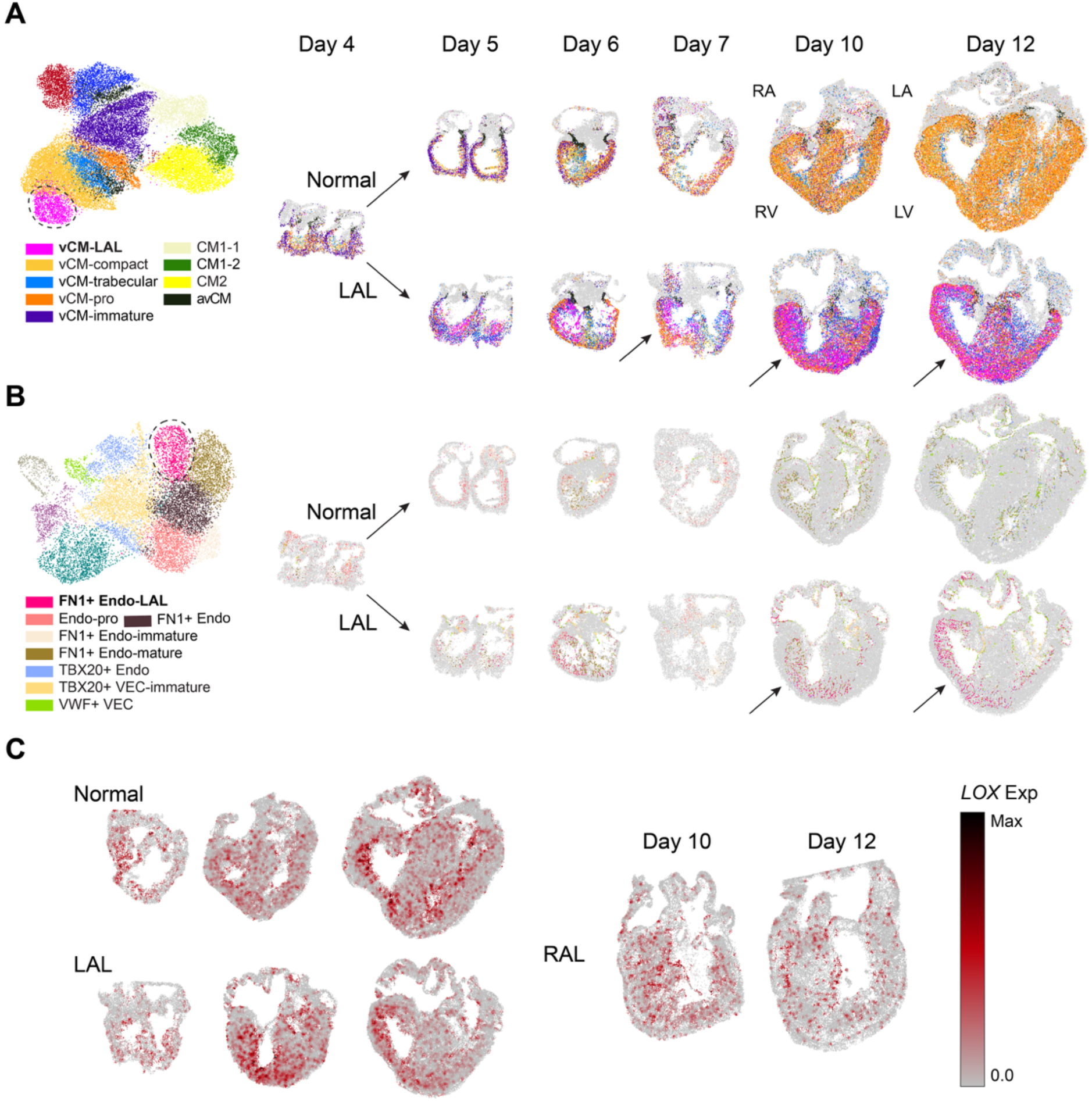
Identification of LOX expressing LAL-enriched cell types and Their Preferential Localization in the RV. (A) UMAP plot of cardiomyocytes from scRNA-seq data, colored by cell type. The vCM-LAL cluster is outlined with a dashed line (left). Spatial maps of vCMs in normal (top right) and LAL (bottom right) hearts across developmental time points. (B) UMAP plot of endothelial cells from scRNA-seq data, colored by cell type. The FN1⁺ Endo-LAL cluster is outlined with a dashed line (left). Spatial maps of non-CECs in normal (top right) and LAL (bottom right) hearts across developmental time points. (C) Spatial expression of *LOX* in normal (left top), LAL (left bottom), and RAL (right) heart across collected developmental time points.

Spatial mapping revealed that these LAL-enriched populations preferentially localize to the over-loaded right ventricle (RV) in LAL hearts (**Figure 3A, 3B, S3**). Notably, *LOX* expression was concentrated in the RV in LAL hearts, whereas in other conditions, *LOX* expression was less pronounced or more evenly distributed across both ventricles (**Figure 3C**). This observation was further supported by LV vs RV differential expression analysis within LAL hearts, where *LOX* ranked among the most highly expressed genes in the RV (**Table S2**). Further supporting a distinct association between *LOX* expression and the LAL-enriched populations, spearman correlation analysis between *LOX* expression and proportion of vCM-LAL or FN1+ Endo-LAL per ST spot revealed positive correlation coefficient in LAL hearts, higher than in other conditions (**Figure S4C**). *LOX* plays a critical role in ECM remodeling and fibrosis, and its upregulation has been implicated in fibrotic remodeling and increased tissue stiffness^25–29^. Consistent with this role, over-representation analysis (ORA) of the LAL-enriched populations highlighted pathways related to wound healing and collagen fibril organization (**Table S1**), indicating that these LAL-enriched cell states contribute to fibrotic remodeling in the mechanically overloaded RV of LAL hearts.

### Hemodynamic Redistribution Induces Ventricular Wall and Lumen Remodeling in Vivo

In normal chick hearts, both left and right ventricular free walls develop compact and trabecular domains. The LV wall develops greater compact volume compared to the RV, while the RV develops greater trabecular compartment volume. The RV develops a greater density of trabeculations, while the LV trabeculations are thicker^30^. The RV and LV lumens are largely similar across development^31^. LAL treatment at day 4 results in hemodynamic de-loading of the left AV valve and ventricle and compensatory over-loading of the right AV valve and ventricle^32^. LAL hearts exhibited a reduced LV lumen size, consistent with its underdevelopment following surgery (**Figures 3A, B, spatial map**). RV lumen size in contrast increased considerably. Interestingly, although the lumen size was decreased, the LV wall was not proportionally thinner. The RV lumen in LAL treated hearts increased considerably, and greater than the proportion of RV wall volume (**Figures 3A, B, spatial map**). RAL treatment induced complementary morphological changes in the chick heart. The size of the LV lumen increased while RV decreased, opposite to LAL treatment (**Figure S6**). These results indicate that hemodynamic loading significantly influences left ventricular lumen and myocardial wall volumes, with compensatory changes in the contralateral ventricle.

To further characterize ventricular wall remodeling in higher resolution, we manually annotated the LV and RV of day 10 and day 12 hearts with napari^33^, guided by mapped cell type and anatomical landmarks, ensuring comprehensive representation of the ventricular wall architecture (**Figure 4A**). From the annotated region, we computed each spot’s normalized spatial coordinate along the outer-inner axis, hereafter referred to as “layer position” (**Figure 4A; STAR Methods**).

**Figure 4.**
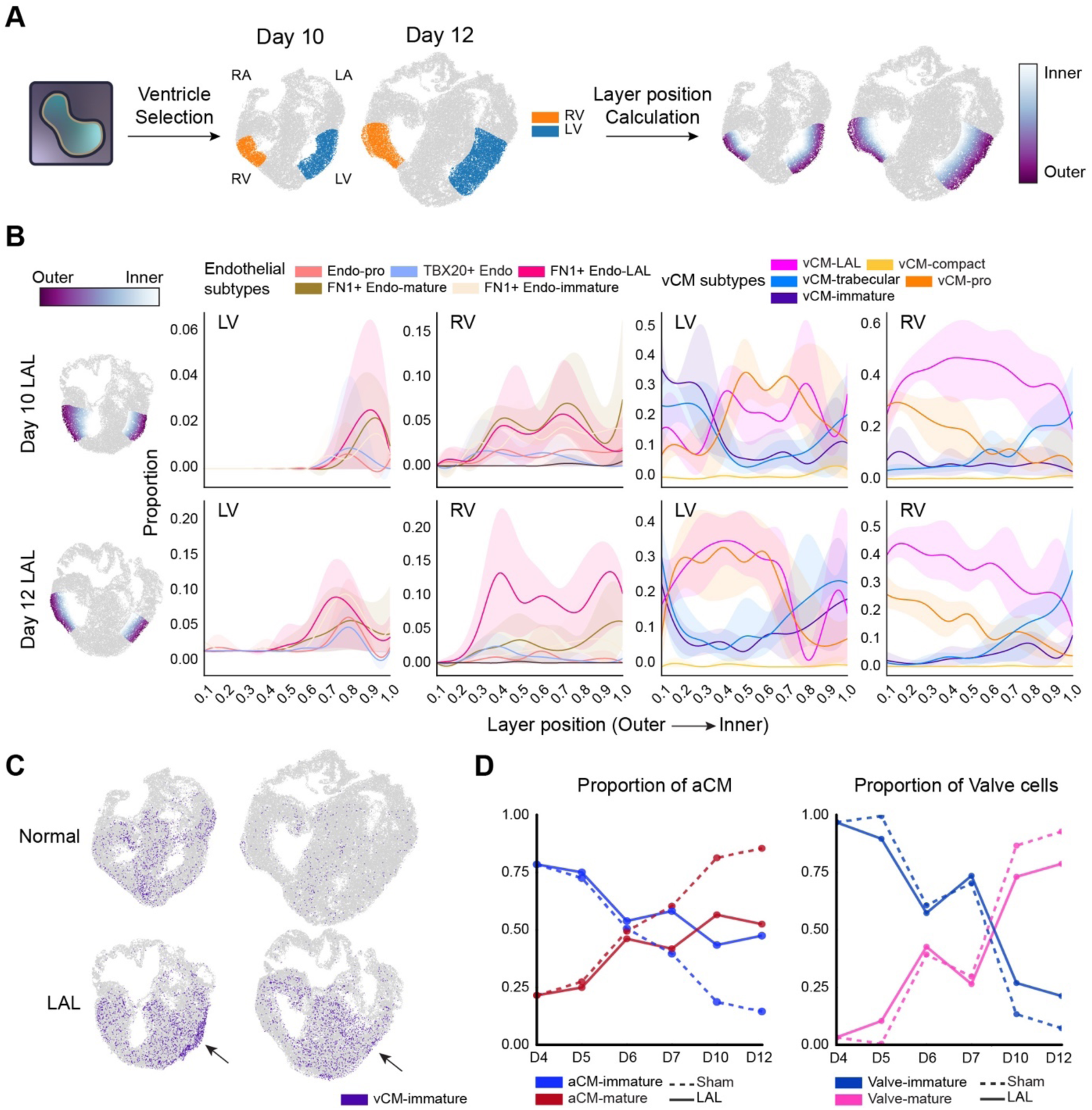
Alteration of Heart Ventricle Architecture and Signs of Lagged Maturation in LAL Heart. (A) Schematic overview of RV and LV annotation and layer position calculation in normal and LAL hearts. (B) Distribution plots showing layer-wise changes in non-vascular endothelial cell and vCM proportions from outer to inner layers of the LV and RV in LAL hearts. (C) Spatial maps of vCM-immature populations at Day 10 and 12 in normal (top) and LAL (bottom) hearts. (D) Line plots of the proportion of aCMs and valve cells across developmental time points (Day 4, 5, 6, 7 and 12) in both normal and LAL conditions.

Both ventricular free walls in normal hearts maintained a balanced cellular architecture, characterized by a distribution of endocardial cells, vCM-trabecular, and vCM-compact proportional to their developed morphology (**Figures 2D, E, S3, and S5**). With LAL treatment, the RV exhibited thickening of the trabecular layer, with endocardial spots extending outward toward the outer layer of the ventricle compared to both the LV of LAL hearts and the RV of normal hearts (**Figure 4B, S5A**). This suggests increased trabeculation leading to trabecular thickening in RV as an adaptive response to the increased volume load following ligation^34–36^. While the RVs of LAL hearts exhibited increased trabeculation, the LVs showed distinct forms of disorganization within the innermost layer. In normal hearts, this layer is primarily composed of endocardial cell subtypes (**Figure S5B**). In contrast, LVs from LAL hearts displayed an aberrant expansion of vCM-trabecular and a persistent presence of vCM-immature population that infiltrated the endocardial layer, potentially leading to functional abnormalities distinct from those observed in the RV (**Figure 4B, S5B**). Trabecular formation is known to be mechanosensitive, and here, our data clearly showcase that different types of cellular disorganization arise in response to either overload or underload of blood flow to the ventricle.

### LAL Hearts Exhibit Signatures of Lagged Maturation

Beyond changes in organization, we also observed signs of delayed maturation in LAL heart development across multiple lineages. In the ventricle, a population of vCM-immature was found localized in the LV of LAL hearts at day 10 and 12, particularly within the outer layer (**Figure 4C, B, S3, S5**). This pattern was absent in the normal heart, where by day 12, vCM-immature was nearly depleted (**Figure 4C**). Furthermore, these lagged maturation signs were found in other areas of heart. While the transition from aCM-immature to aCM-mature typically occurs by day 6 in normal development, in LAL hearts, this shift was delayed until between day 7 and 10, with persistent enrichment of the aCM-immature population during later developmental stages (**Figure 4D, S3**). Transition timing was similar for the valve lineage in normal and LAL. However, LAL hearts exhibited a higher proportion of Valve-immature compared to normal hearts (**Figure 4D, S3**).

A previous study reported that the hemodynamic disturbance induced by RAL was less pronounced than that caused by LAL, and subsequent cardiogenesis was not significantly affected^7^. Consistent with previous findings, we observed the expected loss of the right ventricular lumen, but overall, the cellular dynamics of RAL hearts closely resembled those of normal hearts. RAL hearts retained abundant mature-stage populations, including aCM-mature, Valve-mature, endocardial populations, VECs, CECs, and pericytes localized within the ventricular region (**Figure S6**). However, we observed abundant vCM-trabecular in RAL in scRNA-seq analysis, and spatial mapping confirmed its abundance within the heart ventricle without evidence of spatial preference across regions (**Figure S1, S4A, S6**). This finding suggests that RAL-induced hemodynamic changes may selectively modulate specific cardiomyocyte populations.

### Altered Molecular Environment Reveals Pathological Defects in LAL Ventricles

The alterations in the LAL ventricle organization and retention of immature cells in RV led us to further investigate the molecular consequences of this perturbation. Using manually annotated ventricle regions, we performed DEG analysis followed by gene set enrichment analysis (GSEA) based on Gene Ontology Biological Process (GO BP) gene sets (**Figure 4A, S7 and Table S3, S4**). In normal LV tissue at both day 10 and 12, GSEA revealed significant enrichment of gene sets related to normal cardiac development, supporting the molecular integrity of normal ventricular maturation (**Figure S7 and Table S3**). Day 10 LAL LV exhibited significant enrichment of gene sets indicating molecular adaptation to the altered hemodynamics. Specifically, pathways involved in aerobic respiration and ATP synthesis were upregulated. Interestingly, these same pathways were markedly depleted by day 12, with overlap among leading-edge genes (**Figure 5A** **and Table S3**). This temporal pattern indicates a transient metabolic activation phase characterized by increased oxidative phosphorylation and energy production. Additionally, day 10 LAL LV showed enrichment of broader mitochondrial pathways, reinforcing a temporary shift in cellular metabolism toward energy production (**Figure 5B** **and Table S3**). Comparable dynamics have been reported in early cardiac hypertrophy models, where an initial surge in aerobic metabolism is followed by suppression during maladaptive remodeling^37^. While earlier studies emphasized hypoxia-driven anaerobic metabolism^38,39^, our findings reveal a distinct, time-limited upregulation of energy production, captured through fine-grained temporal sampling, highlighting a previously underappreciated aspect of cardiac adaptation to altered blood flow. Concurrently, we observed increased enrichment of hydrogen peroxide metabolism pathways, which is noteworthy given the dual role of ROS as both a byproduct of mitochondrial respiration and a driver of cardiac injury^40^.

**Figure 5.**
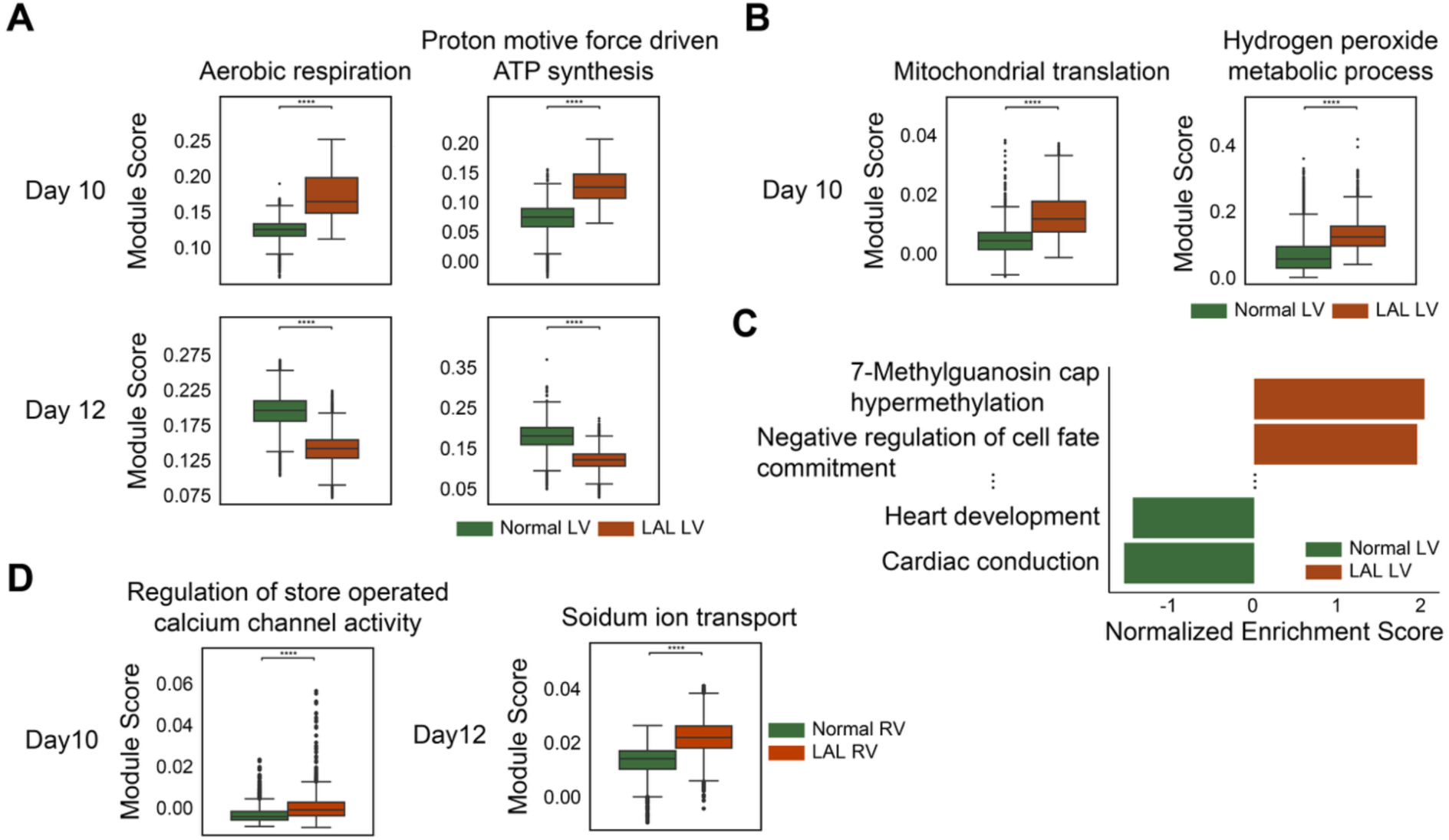
Molecular characterization of LV and RV in normal and LAL Heart. (A) Boxplots of module scores for “Aerobic respiration” and “Proton motive force-driven ATP synthesis” at Day 10 and 12 LVs. Overlapping leading-edge genes from both time points were used for module score calculation. Full GSEA results are available in Table S3. (B) Boxplots of module scores for “Mitochondrial translation” and “Hydrogen peroxide metabolic process” at Day 10 LVs. Full GSEA results are available in Table S3. (C) Bar plot of normalized enrichment scores (NES) showing the top two gene sets enriched in LVs and two gene sets associated with normal heart function enriched in normal LVs at Day 12. Full GSEA result available in Table S3. (D) Boxplots of module scores for “Regulation of store operated calcium channel activity” on Day 10 RVs and “Sodium ion transport” on Day 12 RVs. Full GSEA results are available in **Table S3**.

On day 12, GSEA of the LAL LV revealed significant enrichment of four distinct pathways (**Table S3**). Among these, the second most enriched pathway was negative regulation of cell fate commitment, characterized by five leading-edge genes: *FZD7*, *SFRP2*, *SOSTDC1*, *NKX6-2*, and *LOXL3* (**Figure 5C** **and Tables S3, S4**). Together with the earlier observation of retained ventricular cardiomyocytes exhibiting immature phenotypes (vCM-immature) in the late-stage LAL heart (**Figure 4C**), this enrichment indicates a persistent immature molecular environment that may contribute to such retention.

The RV of LAL hearts at day 10 also showed enrichment of the hydrogen peroxide metabolic process pathway, similar to the LV, although transient changes in aerobic respiration were not observed in the RV (**Table S3**). Interestingly, we observed enrichment of genes related to the regulation of store-operated calcium channel activity, with leading-edge genes *STIM1, STIM2,* and *UBQLN1* (**Figure 5D** **and Table S3, S4**). Previous studies have shown that store-operated Ca²⁺ entry (SOCE) is essential for the development of pathological cardiac hypertrophy, which represents a primary cardiac response to mechanical overload. Moreover, *STIM1* knockdown experiments have demonstrated a complete inhibition of hypertrophic growth in neonatal cardiomyocytes^41–44^. However, these prior studies were conducted in-vitro, and the in-vivo role of store-operated Ca²⁺ entry had not been validated. Given that the LAL RV experiences mechanical overload due to the surgical intervention, the enrichment of this gene set not only highlights its relevance to the overloaded condition but also provides in-vivo evidence supporting its involvement in cardiac hypertrophic responses.

In the day 12 LAL RV, the gene set associated with sodium ion (Na⁺) transport was enriched (**Figure 5E** **and Table S3**). On day 10, we observed increased store-operated calcium channel activity in the LAL RV. Earlier studies have shown that SOCE is closely linked to the regulation of cytosolic Na⁺ levels^45–48^. Therefore, the enrichment of Na⁺ transport likely represents a sustained response to the earlier increase in store-operated Ca²⁺ activity. In addition, several gene sets related to morphogenesis, proliferation, and regeneration, including heart trabeculae, were also enriched (**Table S3**). This finding is consistent with the previously observed thickening of the trabecular layer, providing molecular evidence supporting an adaptive response to mechanical overload (**Figure 4B**).

### Neighbor Population Dynamics in Valve Reveal Plasticity- and Time-Dependent Changes, showing Disrupted Spatiotemporal Cell Neighbors in LAL Valve

Development of heart valves involves complex interactions with their microenvironments, and disturbance of blood-flow homeostasis influences neighboring populations around the heart valve^49^. To investigate the spatial and temporal dynamics of cell populations during heart valve development, we focused on the valve region and applied GraphST, a tool that identifies spatial domains by integrating spatial location and gene expression data^50^. By restricting the analysis to the valve region, we identified spatial subdomains. For each subdomain, we calculated a regional plasticity score and quantified the proportions of neighboring cell types, enabling analysis of plasticity-dependent changes in the cellular microenvironment (**Figure 6A; STAR Methods**). The regional plasticity score was defined as the mean spatial plasticity score of fibroblasts across all spots within each sub-domain, where the spatial plasticity score of fibroblasts was calculated by weighting the plasticity score of each fibroblast subtype (from scRNA-seq velocity analysis) by its proportion within each spot (**Figure S1; STAR Methods**). Higher scores indicate greater potential for cell state transition, whereas lower scores reflect more stable sub-domains, corresponding to the yellow and blue regions in the spatial plots colored by plasticity score (**Figure 6B, 6C**).

**Figure 6.**
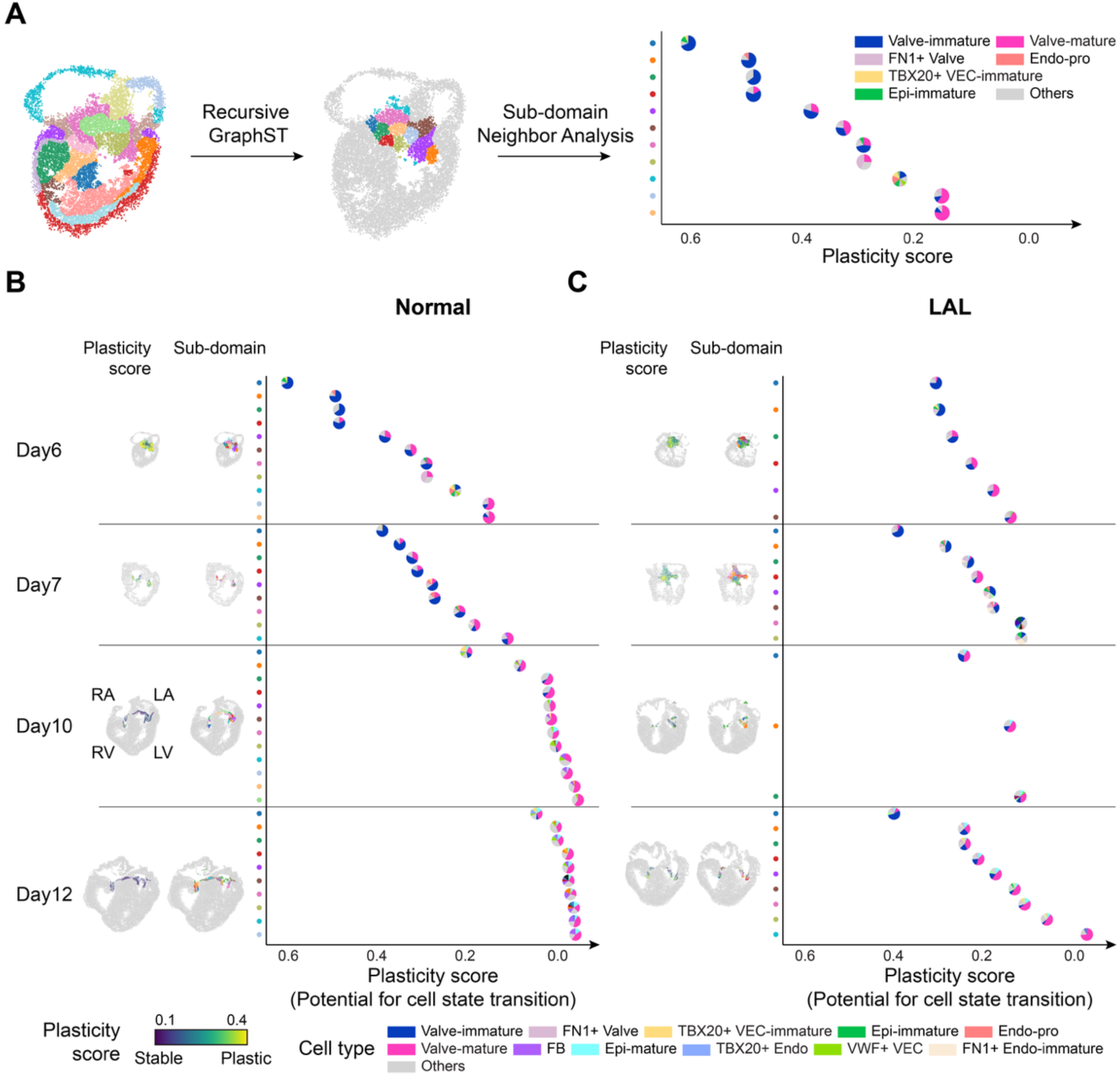
Neighborhood Analysis of Valve Region Revealing Plasticity in LAL Valve. (A) Schematic overview of the neighborhood analysis in valves. Recursive GraphST was applied to define the valve region and its sub-domains. For each sub-domain, plasticity scores were calculated alongside cell type composition. Pie charts represent sub-domains, with slices colored according to cell type composition and positioned according to the plasticity score of the corresponding sub-domain. (B) Normal heart neighborhood dynamics of the valve region across developmental time points (Day 6, 7, 10, and 12). From left to right: spatial plots colored by plasticity score; spatial plots colored by sub-domains; dots colored by sub-domains; and pie charts for each sub-domain colored by cell type composition, positioned according to the plasticity score of each sub-domain. (C) LAL heart neighborhood dynamics of the valve region across developmental time points (Day 6, 7, 10, and 12). From left to right: spatial plots colored by plasticity score; spatial plots colored by sub-domains; dots colored by sub-domains; and pie charts for each sub-domain colored by cell type composition, positioned according to the plasticity score of each sub-domain.

In both normal and LAL heart valves, we observed that highly plastic domains were predominantly localized to the distal tips of the valve region (**Figure 6B, 6C spatial plots**). In the normal heart, temporal alignment of sub-domains across developmental time points revealed a progressive rightward shift, consistent with a gradual and spatially organized maturation of the valve structure (**Figure 6B**). In contrast, this directional shift was absent in the LAL heart, suggesting that the LAL-induced shift in pressure and normal forces disrupts the spatiotemporal progression of valve maturation (**Figure 6C**).

Neighborhood analysis of sub-domains further revealed distinct differences in cell neighbors between normal and LAL valves. In the normal valves, plastic/early domains were surrounded by developmentally immature cell types such as TBX20⁺ VEC-immature, Epi-immature, and proliferative Endo-pro cells (**Figure 6B**). In contrast, stable/late neighborhoods in the normal heart were composed of more mature and structurally supportive cell types such as TBX20+ Endo, Epi-mature, and VWF+ VEC. TBX20+ Endo expresses key components of ECM, such as type I and III fibrillar collagens and maintains TBX20 (**Table S1**). Notably, fibrillar collagens are major components of the developing valve matrix, and TBX20 is critical for valve endocardial cell proliferation and ECM organization^51–54^. Epi-mature expressed both type III collagen and elastin, contributing to the valve’s elastic properties^55^. VWF+ VEC, a vascular endothelial cell type, emerged later in development and localized near the valve (**Figure 6B, 2E**).

In the LAL heart valves, although the constituent cell types were largely preserved, their spatial organization across plastic to stable domains was disrupted. TBX20+ VEC-immature, typically restricted to early-phase neighborhoods, was aberrantly observed in stable/late domains at day 10 and 12 (**Figure 6B, C**). Conversely, Epi-mature, normally appearing from day 10 as a late-phase neighbor, appeared prematurely at day 7 (**Figure 6B, C**). Furthermore, these two temporally segregated populations were co-localized in LAL valves, whereas they remained clearly separated in the normal heart. FN1+ Endo-immature, typically a minor neighbor at day 7 in normal valves, occupied a larger proportion in LAL (**Figure 6B, C**). Collectively, these observations point to a breakdown in the spatial and temporal fidelity of cellular neighborhood organization in LAL heart valves, consistent with delayed progression observed in the spatial cell type maps of the LAL heart, which is consistent with previous finding (**Figure 4D, S3**)^56^.

## DISCUSSION

In this study, we present the first spatiotemporal atlas of heart development under experimentally altered hemodynamics, offering new insights into how mechanical forces shape cellular and molecular programs through unbiased, systematic analysis. By performing left and right atrial ligations (LAL and RAL) in developing hearts, we induced controlled hemodynamic perturbations and collected samples at later developmental stages than prior studies^7–9^. We integrated high-resolution spatial transcriptomic (ST) datasets with paired single-cell transcriptomics to trace the consequences of these perturbations. This dataset enabled characterization of heart development under distinct hemodynamic conditions, identification of local tissue remodeling, detection of LAL-enriched cell types with preferential RV localization, ventricular wall disorganization, and molecular alterations in the ventricle. Together, these findings provided new insights into how biomechanical forces affect heart development across scales.

Our findings are consistent with prior understanding of key morphogenic programs in the heart, including epicardial derivation of the atrioventricular fibrous skeleton, timing of the compact and trabecular ventricular growth and maturation, and maturation of the atrioventricular valves. High resolution spatial transcriptomics maps these diverse lineages and their phenotype transitions in space and time in the same embryo. We further establish spatial and temporal evolution of heterogeneous cellular dynamics as an underrecognized hallmark of these events. While much has been learned about the genetic basis of morphogenesis, genetic perturbations are less suited to interrogate later-stage morphogenesis where multiple cell types collaborate to form and mature tissues. For example, timed ablation of lineages at late stages can yield minimal phenotypes, suggesting that local cellular communities can compensate to sustain morphogenesis. These communities appear highly sensitive to their biomechanical environment, forming and sizing tissue compartments in response to local forces. Malformation at these later stages can therefore reflect acquisition of an alternative morphological state that satisfies local biomechanical cues without requiring genetic mutation. Altogether, our results provide a rich resource for identifying programs operating across diverse cellular communities to progress local cardiac morphogenesis, and similarly how community phenotype divergence associates with malformation decisions.

Our results identify cellular plasticity as an indicator of local tissue maturation. In the atrioventricular valves, maturation progressed from the base towards the tip, whereas in the myocardium the trabecular layer was less plastic than the compact zone, which continues to proliferate over fetal morphogenesis. Proper growth and maturation of these tissue compartments are essential to sustain pumping power and unidirectional circulation while the rest of the growing fetus demands increasing nutrient supply. Retention of plasticity may represent a cellular adaptation to altered biomechanics that precedes morphological change. Our results suggest that plasticity is a local rather than an individual cell phenotype characteristic, supporting the idea that heterogeneous cellular communities respond collectively to microenvironmental change. We speculate that a reduction in plasticity could mark a local restriction in adaptation to microenvironmental perturbation, potentially signaling a transition towards pathogenic remodeling. In our results, the endocardium stabilized earlier in LAL treatment relative to normal. The emergence of abundant stable *LOX* expressing FN1+ Endo-LAL and vCM-LAL cells with LAL is consistent with a transition in RV remodeling from adaptive to pathological in response to overload, and similarly with the disruption of endocardial boundary in the LV in response to under-loading. These features may have relevance for decoding the pathogenesis of congenital malformations. Delayed maturation in LAL hearts was previously attributed to reduced ventricular volume or altered shear stress^7,8,32,57,58^. Here, we demonstrate the delayed maturation through the persistence of immature cells at later stages and the enriched negative regulation of cell fate commitment. This highlights that altered hemodynamics not only remodel heart structure but also maintains cellular immaturity.

The chick embryo is an experimentally robust and well characterized model system. Its bi-ventricular composition and electrophysiology is closer to human than the mouse, and its larger size at similar stages provides greater structural complexity than the fish or mouse, more similar to human^31^. Experiments to date affirm conserved cell lineages, molecular regulation and cellular differentiation between the chick and mammalian systems^59–65^. While genetic perturbation is possible but more difficult in the chick than the mouse or fish, the chick exhibits unparalleled advantages in experimental manipulation to test mechanisms of mechanics and hemodynamics in clinically relevant contexts^66–68^. Mechanical perturbation in live fetal mice is not possible, and iPS-cell derived organoids are too crude to test these hypotheses^69^. Eggs enable high experimental throughput with no collateral loss of life critical to test mechanobiological hypotheses. We recapitulate key features of hemodynamic signaling in ventricular malformation: partial LAL constricts inflow and unloads the LV, resulting in a stunted chamber lumen, thickened myocardial wall with reduced trabeculation, and, in advanced stages, endomyocardial fibroelastosis, which are classic features of human HLHS^66,70.^

In addition to this experimental design, we applied a computational strategy to combine cell-state dynamics inferred from scRNA-seq data with spatial transcriptomics for studying heart development. RNA velocity analysis, which infers transcriptional dynamics from the ratio of spliced and unspliced mRNA counts, has been widely applied in scRNA-seq studies. More recently, several studies have extended this approach to spatial transcriptomics^71–74^. However, existing approaches have limited capacity to resolve cell type-specific dynamics or to transfer the dynamic information derived from scRNA-seq into spatial coordinates. To capture cell-state transitions during cardiac development at cell-type resolution, we introduce a plasticity score that quantifies both the magnitude and probability of cell-state transitions^10^ (**STAR Methods**). Given that spatial transcriptomics achieves near single-cell but not true single-cell resolution, we projected these plasticity scores onto spatial data by weighting them according to the probabilistic cell-type composition of each spatial spot. This integration enables the visualization and analysis of active cell-state transitions within their anatomical context, providing a powerful approach to investigate dynamic cellular behaviors during tissue morphogenesis.

Tissue morphogenesis emerges from dynamic interactions between diverse cell types. Previous studies of the cardiac microenvironment have relied on *in vitro* co-culture systems^75–82^. While informative, these approaches often require extensive manipulation, such as tissue isolation or reconstruction of artificial environments. Similarly, single-cell and single-nucleus RNA sequencing lack spatial context and cannot fully capture cellular communities within intact tissue. Spatial transcriptomics overcomes these limitations by enabling unbiased analysis while preserving the spatial architecture of the developing heart. Our analysis demonstrates that ST can resolve stage-specific microenvironments and perturbation-associated alterations directly within native tissue context.

Yet, while spatiotemporal transcriptomic analysis captures key cellular and molecular dynamics, there is still room for finer characterization. Here, we used Seeker (Curio) which provides near single-cell level resolution, so each spot may contain transcripts from more than one cell, which is a current limitation of most spatial platforms. Further study using true single-cell-resolution approaches will enable finer characterization and may reveal structures that are currently missed, for example, neural cells detectable by scRNA-seq but not confidently mapped in ST.

Despite these inherent limitations, our analysis revealed several robust and biologically meaningful findings, providing comprehensive, high-resolution insights into cellular and molecular dynamics under different hemodynamic conditions and illuminating how hemodynamic stress disrupts heart development. Our study also provides a generalizable framework to investigate the complex interplay between mechanobiology and cellular and molecular programs during heart development.

## STAR METHODS

### Heart Ligation

Fertilized *Bovans Brown* chicken eggs were incubated in a forced-air incubator at 37.5 °C and 50% relative humidity until reaching Hamburger-Hamilton stage 24 (HH24) at approximately 4.5 days of development. At HH24, the eggs were positioned with the air cell facing upward, and the shell overlying the air cell was carefully cracked and removed using dissection forceps to expose the air sac. The inner shell membrane was gently removed to expose the embryo while minimizing disruption and avoiding bleeding. At this stage, the embryo typically lies on its left lateral side, allowing visualization of the right atrium for right atrial ligation; to perform left atrial ligation, embryos were gently rotated to expose the left atrium. Following orientation, the amnion surrounding the heart was carefully opened using fine forceps to expose the beating heart. A pre-tied 10-0 nylon suture loop (10-0 Pocket Suture™ Black Nylon Micro Monofilament Training Sutures, TS1012) with a loop diameter of approximately 1 mm was positioned around the target atrium, and the free ends of the suture were tightened using fine forceps to constrict the atrium, after which excess suture was trimmed using micro-scissors. Embryos undergoing right atrial ligation remained in their original orientation, whereas those undergoing left atrial ligation were rotated back to their normal position following the procedure. After ligation, the windowed eggs were sealed using Saniderm® medical polyurethane film dressing to provide a sterile, breathable barrier and prevent contamination during continued incubation.

### Sample Preparation for Single-cell RNA Sequencing

We used fertile bovans brown chicken (Gallus Gallus) eggs to study the development under normal conditions on HH24, 26, 31 and 36 (translated to day 4, 5, 7, and 10 for simplicity). For development under hemodynamic perturbation, either the left or the right atria were ligated after day 4 for LAL and RAL and hearts were collected at subsequent time points. Hearts were in ice-cold Hank’s Balanced Salt Solution (HBSS). For day 4 normal time points, 13 whole hearts were used. For day 5 normal, LAL and RAL, 8 hearts each i.e 24 hearts total were used. For day 7 it was 2 hearts for each condition (6 total) and for day 10, it was one heart each (3 total). Cells obtained using this protocol tested for a high RNA integrity number and were passed further for fixation. Day 4, 5 and 7 samples were digested using 300 U/mL of collagenase type II for one cycle of 15 mins under mild agitation at 37 C. The cells were spun down and digested again for 10 mins. For day 10 hearts, the organs were chopped and then incubated with collagenase for 20 min and 10 mins. These cells were passed through a 40 um filter, pelleted and resuspended in a red blood cell lysis buffer. Cells were resuspended in a phosphate-buffered saline solution. Cells were counted using an automated hemocytometer and prepped for fixation using the Parse evercode cell fixation v2. Post-fixation, the cells were stored at -80 C. The fixed cells from all 10 samples were processed using Parse evercode WT v2 kit designed for 100,000 cells. The 8 final sub-libraries were run on a single lane using the illumina P3 flow cell 100 bp kit with configuration 46+6+86 (cDNA + i7 index + Barcoding). Gallus gallus genome and annotations (assembly: GRCg7b maternal broiler) were downloaded from Ensembl genome browser and processed using Parse bioscience computational pipeline v1.0.4.

### Preprocessing of scRNA-seq data

Preprocessing of scRNA-seq data was performed using the scanpy package^83^. Cells were filtered out based on the following criteria: (1) doublet score > 0.25, calculated using the scrublet package^84^, (2) fewer than 300 genes or more than 10,000 genes, and (3) mitochondrial molecule fraction > 10%, yielding 80,197 cells with 28,140 genes, with median transcripts/ cell of 1,503, median gene/cell of 931.

The raw count matrix was then normalized by (1) feature counts divided by total feature counts in each cell, scaled by 10,000, and subjected to natural-log transformation with a pseudo-count of 1 (sc.pp.log1p). The features were then scaled and centered. Principal component analysis (PCA) was performed on the 2,000 highly variable genes (HVG), and the top 30 PCs were selected based on the knee-point in the elbow plot.

### Cell type Annotation

#### Coarse Cell type Annotation

We used hierarchical strategy for cell type annotation, beginning with coarse resolution (9 major cell types) followed by granular subtyping. For the coarse resolution, a *KNN* graph was constructed (k=15) on the Euclidean distance in PCA space based on the top 30 PCs. Leiden clustering was applied with resolution=0.7, yielding 21 clusters. Clusters were annotated based on the expression of canonical markers genes, including *TBX18, ALDH1A2, WT1, TGFB2, and TCF21* for epithelial cells, *COL1A1, PDGFRA, and COL3A1* for fibroblasts, *PECAM1*, *CD34*, and *CDH5* for endothelial cells, *TNNT2*, *MYH7*, *ACTN2*, *TTN*, *ACTC1* for cardiomyocytes, *HBBA*, *HBAD*, and *HBA1* for Erythrocytes, *PTPRC MERTK*, and *CSF1R* for macrophages, *MYH11*, *TAGLN*, and *ACTA2* for Mural cells, *CELF4*, and *MAP2* for Neural cells, *PTH*, *GCM2* for Parathyroid cells (**Figure S1**).

#### Granular Cell type Annotation of Epithelial cells, Fibroblasts, Cardiomyocytes and Endothelial cells

Following coarse cell type annotation, epithelial cells, fibroblasts, cardiomyocytes and endothelial cells were subsetted. Each subset was re-scaled, and 2,000 HVGs were identified respectively. PCA was performed on the subset-specific HVGs, and the top 20 PCs were selected using the knee-point. Leiden clustering was applied with resolution=0.7 for cardiomyocytes and endothelial cells, resolution=0.6 for epithelial cells, and resolution=0.5 for fibroblasts. Subtype annotation was performed using known marker genes and differentially expressed genes (DEGs), identified with scanpy ‘sc.tl.rank_genes_groups’ function (**Figure S1 and Table S1**).

### Sample Preparation for Slide-seq spatial transcriptomics platform

First, the heart was carefully harvested and placed in a 10mm Petri dish filled with enough HBSS (Gibco 14025076) to completely submerge it. A 10 ml syringe filled with HBSS and a 33G syringe needle was used, and all air bubbles were expelled before the needle was inserted into the heart through the apex. The syringe plunger was then gently pressed to flush out any remaining blood. Next, a metal freezing dish large enough to hold the entire heart was prepared, and it was half-filled with OCT (Optimal Cutting Temperature compound). The heart was placed in the dish and gently moved around in the OCT to disperse any residual HBSS, then the remainder of the dish was filled with OCT until it was full. The heart was positioned in the center of the dish with the outflow tracts facing upwards and was gently pressed against the bottom of the dish to ensure it stayed in place. 2-Methylbutane was poured into a plastic container, making sure that the liquid level didn’t exceed the height of the freezing dish, and the container was placed into liquid nitrogen until the 2-Methylbutane was completely frozen. Once frozen, the freezing dish was set on top of the solid 2-Methylbutane, and the OCT was allowed to fully solidify and appear white. At this point, the solidified OCT block was transferred into a tissue cassette by flipping the freezing dish and giving it a gentle knock to release the block. Finally, the OCT block was stored at -80°C. For timepoints day 4 and 5, hearts in orientation frontal and sagittal were frozen on the same block. For day 6 and 7, individual hearts were frozen in a block. For timepoints day 10 and 12, 4 hearts were frozen in the same block (2: day 10 and 2: day 12). Here, we used Curio Seeker for the spatial transcriptomics experiments. For timepoints day 4, 5, 6 and 7, individual 3×3 tiles were used. For day 10 and 12 hearts, one tile with all 4 hearts (day 10 and 12 for normal and LAL hearts) was used. An additional 10×10 tile was used for RAL day 10 and 12 hearts. The tile consists of 10 um beads, randomly dispersed in a plane. These beads possess DNA oligos that containing a PCR handle, two bead barcode sequences 1 and 2 for x and y coordinates connected via a linker, a unique molecule identifier (UMI), and a 3’ poly(dT) tail to capture mRNAs. These beads are approximately the size of a cells and capture mostly homogeneous transcriptomes from unique cells. Once a 10 um section was placed on the tile, a cDNA was reverse transcribed from the mRNA. The beads were then dissociated and a second strand was synthesized on the cDNA. This cDNA was amplified to create a cDNA library. Some DNA from this library was tagmented and indexed for the subsequent run on a flow lane. Following sequencing, fastqs were mapped on Gallus gallus reference genome GRCg7b with slide_snake which is the pipeline for snakemake^85^.

### Preprocessing of Spatial transcriptomic data

Preprocessing of Spatial transcriptomic data (Seeker, Curio) was performed with the scanpy package^83^. To remove smears, we calculated a pairwise Euclidean distance matrix between all spots and computed the number of neighboring spots within 100 µm radius. Spots with lower than 100 total counts and neighboring spots over 15 were retrieved. To address the inherent sparsity and noise in spatial transcriptomics data, we applied Smoothie, a method that denoises spatial transcriptomics data with Gaussian smoothing^86^. We applied the following parameters: grid_based_or_not=False, gaussian_sd=46.37, and min_spots_under_gaussian=10. The smoothed Slide-seq data were used for all subsequent analyses, except for cell2location, which requires raw counts as an input.

### Cell type Deconvolution

We used cell2location for Slide-seq data deconvolution^11^. Cell2location is a Bayesian probabilistic spatial deconvolution tool that leverages cell type specific gene expression signatures from the single-cell reference to map profiles onto spatial data, estimating abundance and spatial distribution of cell types.

#### Reference Construction with scRNA-seq data

Separate reference models were constructed for Normal heart, LAL heart, and RAL heart to account for differences in cell type abundance and composition under perturbed hemodynamics. It was done by subsetting cells from our matching scAtlas and training the reference model. In all cases, parameters were set to cell_count_cutoff=5, cell_percentage_cutoff2=0.03, nonz_mean_cutoff=1.12.

#### Cell type Deconvolution

Raw counts for each sample were trained on their corresponding condition reference to estimate the cell type abundances, and we converted the abundance to proportion as:

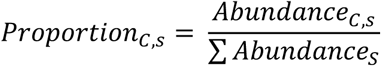

where *C* denotes cell type and *s* denotes spot. From the estimated abundance, each spot was assigned to the cell type with the highest proportion. Despite prior removal of blood contamination, some spots were annotated as erythrocytes, indicating residual blood cells in the tissue. These erythrocyte-assigned spots were excluded from all downstream analyses.

### Velocity Analysis and Calculation of Plasticity Score

#### Velocity Analysis in Epithelial cells, Fibroblasts, Cardiomyocytes, and Endothelial cells

We performed RNA velocity analysis on epithelial cells, fibroblasts, cardiomyocytes, and endothelial cells using scVelo, which infers transcriptional dynamics by modeling splicing kinetics in a likelihood-based framework^10^. The analysis was conducted with nPCs=50 and n_neighbors=30, and the stochastic mode for velocity estimation.

#### Plasticity Score Calculation for Each Cell types

To quantify the degree of transcriptional progression along the inferred velocity, we developed a plasticity score that integrates both the magnitude of velocity and the net flow of transcriptional transitions.

First, the velocity magnitude for each cell *c* was calculated as:

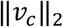

the Euclidean (L2) norm of its velocity vector projected into the UMAP space *v_c_*. This metric was used as a proxy for the speed of change, with higher values corresponding to more dynamic transcriptional states and lower values to more stable states. The values were scaled to the range [0,1] using min-max scaling.

Second, we computed a net flow score from the transition matrix based on the directed transition matrix adata.uns["velocity_graph"]. For each cell, outflow was defined as the sum of transition probabilities in its corresponding row, and inflow as the sum of probabilities in its corresponding column. To avoid bias from unconnected cells, the mean outflow and inflow were computed after excluding zero-valued entries. The net flow score for each cell *c* was calculated as:

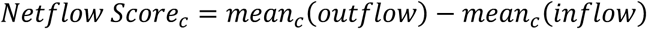

such that values indicate cells acting as sources and lower values indicate sinks. This score was also min-max scaled to [0,1].

The plasticity score for each cell, c, was then defined as the product of its scaled velocity magnitude and scaled net flow score as:

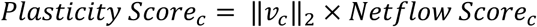

Finally, cell type-level plasticity scores (*Plasticity Scorec*) were obtained by averaging the cell-level plasticity scores for each cell type *C* as:

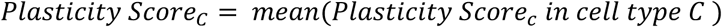

Cell type-level scores were again min-max scaled. Higher scores indicate more plastic cell types, whereas lower scores correspond to more stable cell types.

#### Mapping Plasticity Score on Slide-seq data

To map the plasticity score for each cell type (*Plasticity Score_C_*) in Slide-seq data, we multiplied it to the proportion of cell type *C* in each spot *s* as:

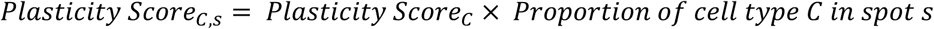

### Neighborhood Analysis in Valve Region

#### Identification of Valve Region Sub-domains with Recursive GraphST

GraphST is a graph-based self-supervised contrastive learning method that uses not only the gene expression profile of each spot but also the spatial coordinates to generate spatially informed domains^50^. To perform high-resolution neighborhood analysis within the valve region, we applied GraphST recursively. In the first round, GraphST was applied to the entire heart tissue to identify the valve domain. In the second round, GraphST was applied only to the selected valve domain. This analysis was performed on smoothed spatial transcriptomics data independently, using ‘mclust’ for clustering, which provides a computationally and time-efficient approach. For the first round, we determined the optimal number of domains by computing the Adjust Random Index (ARI) across different cluster numbers and selecting the range where five consecutive results had ARI > 0.65. Within this range, the final number of clusters was refined through visual inspection. The valve domain was selected based on cell type composition, prioritizing domains with the highest proportion of valve cells. The second round was done the same with the first round, but without visual inspection and with windows of four.

#### Plasticity Score Calculation for Sub-domains

Plasticity scores for each sub-domain were computed as the average of the sum of *Plasticity Score_C,s_* where *C* spans to fibroblasts subtypes. Sub-domains were then ranked from most plastic to most stable and relabeled accordingly, ensuring sub-domain color coding is consistent across different time points and different conditions.

#### Identification of Neighboring Cell types in Subdomains

For each sub-domain, we calculated the proportion of each cell type, which was used to generate the pie chart in **Figure 5**. Cell types were included cumulatively up to 90% of the total composition within each sub-domain, restricting inclusion to those contributing at least 5%. For each sample, a cell type was considered a neighbor if it appeared in at least two sub-domains and further shared across time points.

### Endothelial and vCM Layer Analysis in Heart Ventricle

#### Manual Annotation of Left Ventricle with Napari

Napari is an interactive Python tool for visualizing multi-dimensional images and allows manual selection and annotation of regions of interest^33^. We used Napari to manually select the Left Ventricle (LV) and Right Ventricle (RV) from day 10 and 12 heart from both normal and LAL samples. Selection was guided by knowledge of anatomy of the developing chick heart together with cell type annotations for each spot. We ensured we captured the full ventricular wall, from the epicardial to the endocardial layer.

#### Layer position Calculation

To quantify the spatial position of each spot within the ventricular wall, we calculated a normalized “Layer position” based on the relative distance of each spot to manually annotated inner and outer boundaries of the ventricle. The inner and outer boundary from the annotated ventricle where the boundary is set as a single layer of spots that are in the innermost and outermost area. Then the Euclidean distance from each spot to the nearest inner and outer boundary was then computed. A normalized layer position was defined as:

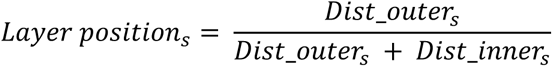

This normalization scales each spot’s position from 0 (outer) to 1 (inner), effectively capturing the transmural gradient across the ventricular wall.

An analogous procedure was applied using the upper and bottom boundary, selecting a single layer of spots that are in the uppermost and bottommost area. This upper-bottom layer positioning was used to confirm that the distribution pattern is consistently maintained across different top-to-bottom regions of the ventricle.

#### Quantification of Endothelial and vCM Distribution Across Layers

In order to see the simple trend of endothelial and vCM distribution, we binned the continuous “layer position” into 10 bins. To capture the regional variation along the ventricular length, the “upper-bottom position” into 5 bins.

Within each upper-bottom bin (# bin= 5), we computed the proportion of each vCM or endothelial cell subtype per layer position bin (# bin = 10) as:

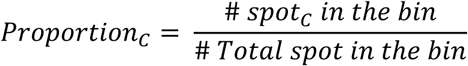

where # spot_C_ is the number of spots predicted as that subtype divided by the total number of spots in the bin. These fractional profiles were averaged across all upper-bottom segments to obtain a mean distribution profile for each subtype. To quantify variability, we calculated the 95% CI using the 2.5th and 97.5th percentiles of the fractions across segments which is indicated as a shade region in **Figure 4** **and S5**.

### Differential Expression and Pathway Analysis

#### Identifying DEGs

Within napari annotated LV regions, we performed Differentially Expressed Gene (DEG) analysis using the pre-built function ‘sc.tl.rank_genes_groups’ from scanpy. Genes with an adjusted p-value less than 0.05 were considered significantly differentially expressed.

#### Gene Set Enrichment Analysis

We conducted Gene Set Enrichment Analysis (GSEA) using the fGSEA package on pre-ranked genes^87^. Ranking was based on the ‘score’ output from sc.tl.rank_genes_groups, with 10,000 permutations performed. The Gene Ontology Biological Process (GOBP) gene sets were used as the reference. Gene sets with an adjusted p-value less than 0.05 were considered significantly enriched. To summarize multiple significant gene sets, when more than five were identified, we calculated the semantic similarity matrix of GO terms using the ‘GO_similarity’ function and clustered these terms with ‘simplifyGO’ from the simplifyEnrichment package^88^. Representative GO terms for each cluster were then selected using ‘reduceSimMatrix’ from the rrvgo package^89^. Module score was calculated based on the leading-edge genes, with the function ‘sc.tl.score_genes’ from scanpy.

#### Over-representation Analysis

We performed Over-Representation Analysis (ORA) using the clusterProfiler package on genes with high expression (log₂ fold change > 0.5 and adjusted p-value < 0.05) using GOBP gene sets^90^. The analysis parameters were set as follows: *pvalueCutoff = 1*, *qvalueCutoff = 1*, and *minGSSize = 1*. Only significant results (adjusted p-value < 0.05) were considered, and gene sets containing the LOX gene were further highlighted (**Table S1**).

## DATA AVAILABILITY

All raw data and analysis-ready datasets are available at Gene Expression Omnibus repository: GSE311835

## CODE AVAILABILITY

The data analysis scripts have been made available on the GitHub repository (https://github.com/parkjooyoung99/Chicken-Heart-Hemodynamics)

## ACKNOWLEDGEMENTS

We acknowledge funding from the National Institutes of Health (HL160028), the National Science Foundation EF-2222434, and the Additional Ventures Foundation. Support was also provided by seed grants from the Center for Vertebrate Genomics at Cornell University.

## AUTHOR CONTRIBUTIONS

JP, SS, RA, IDV, and JB conceived of the study. SS, PL, and HM performed the animal experiments. PL, HM, RA, SS performed single-cell experiments. SS, MLSP, and RA performed the spatial transcriptomics experiments. JP, RA, and AS performed the data analysis. JP, JB and IDV interpreted results. JP, JB and IDV wrote the manuscript. All authors provided feedback and comments.

## SUPPLEMENTAL INFORMATION

Table S1. Differentially expressed genes for each cell types and over-representation analysis for LAL-enriched cell types

Table S2. Differentially expressed genes for LV and RV in day 10 and 12 LAL heart Table S3. Gene set enrichment analysis results, related Figure 5

Table S4. Differentially expressed genes comparing normal and LAL heart LV or RV in day 10 and 12, related Figure 5

---

**Figure S1.**
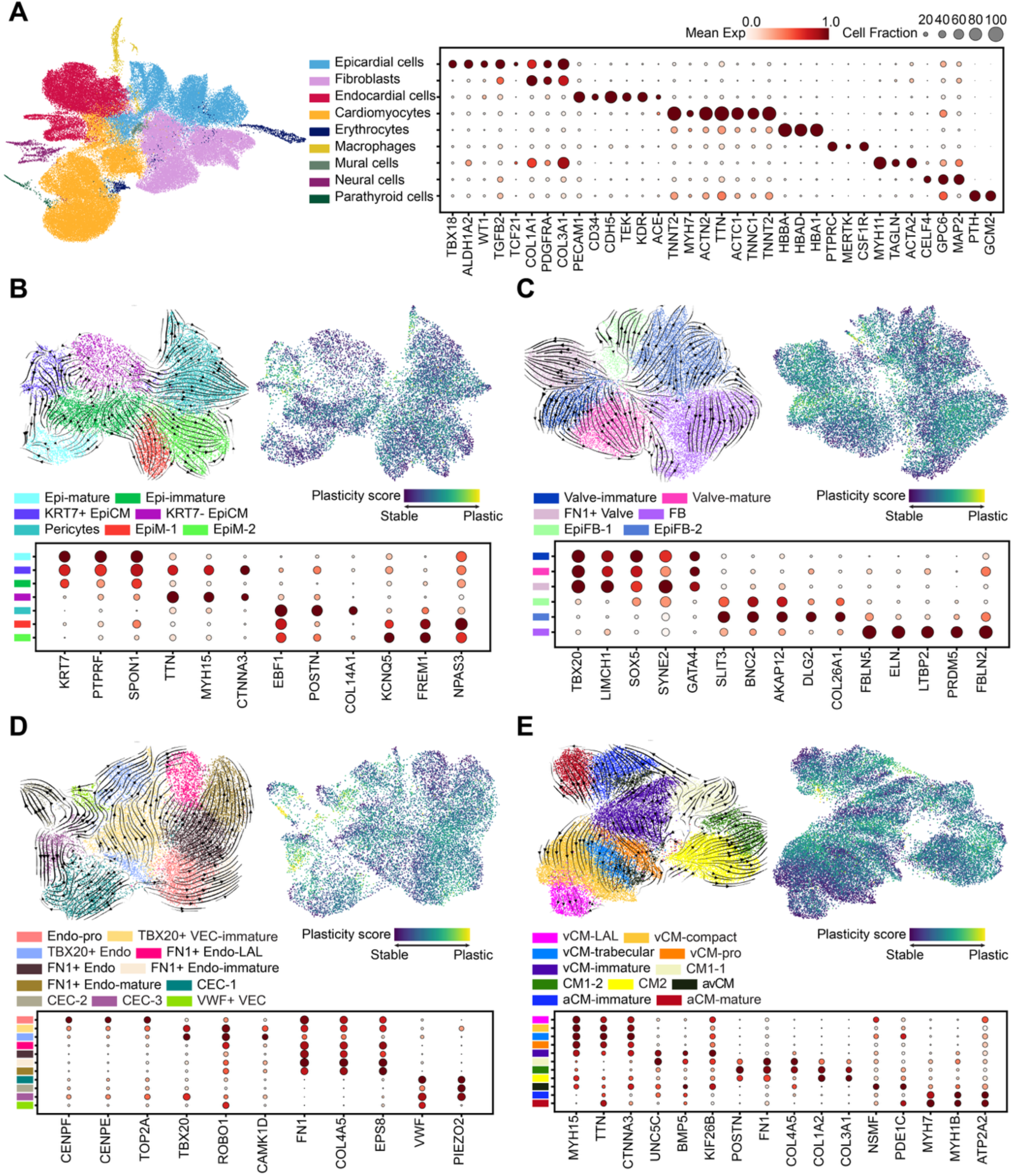
Cell type Annotation in scRNA-seq. (A) UMAP plot of global cell type annotation (left) and dot plot of marker genes for each cell type (right). (B) Subtype annotation of epithelial cells. UMAP plot of epithelial subtypes (left), plasticity scores overlaid with velocity embeddings (right), and dot plot of subtype marker genes. (C) Subtype annotation of fibroblasts. UMAP plot of fibroblast subtypes (left), plasticity scores with velocity embeddings (right), and dot plot of marker genes. (D) Subtype annotation of endothelial cells. UMAP plot of endothelial subtypes (left), plasticity scores with velocity embeddings (right), and dot plot of marker genes. (E) Subtype annotation of cardiomyocytes. UMAP plot of cardiomyocyte subtypes (left), plasticity scores with velocity embeddings (right), and dot plot of marker genes.

**Figure S2.**
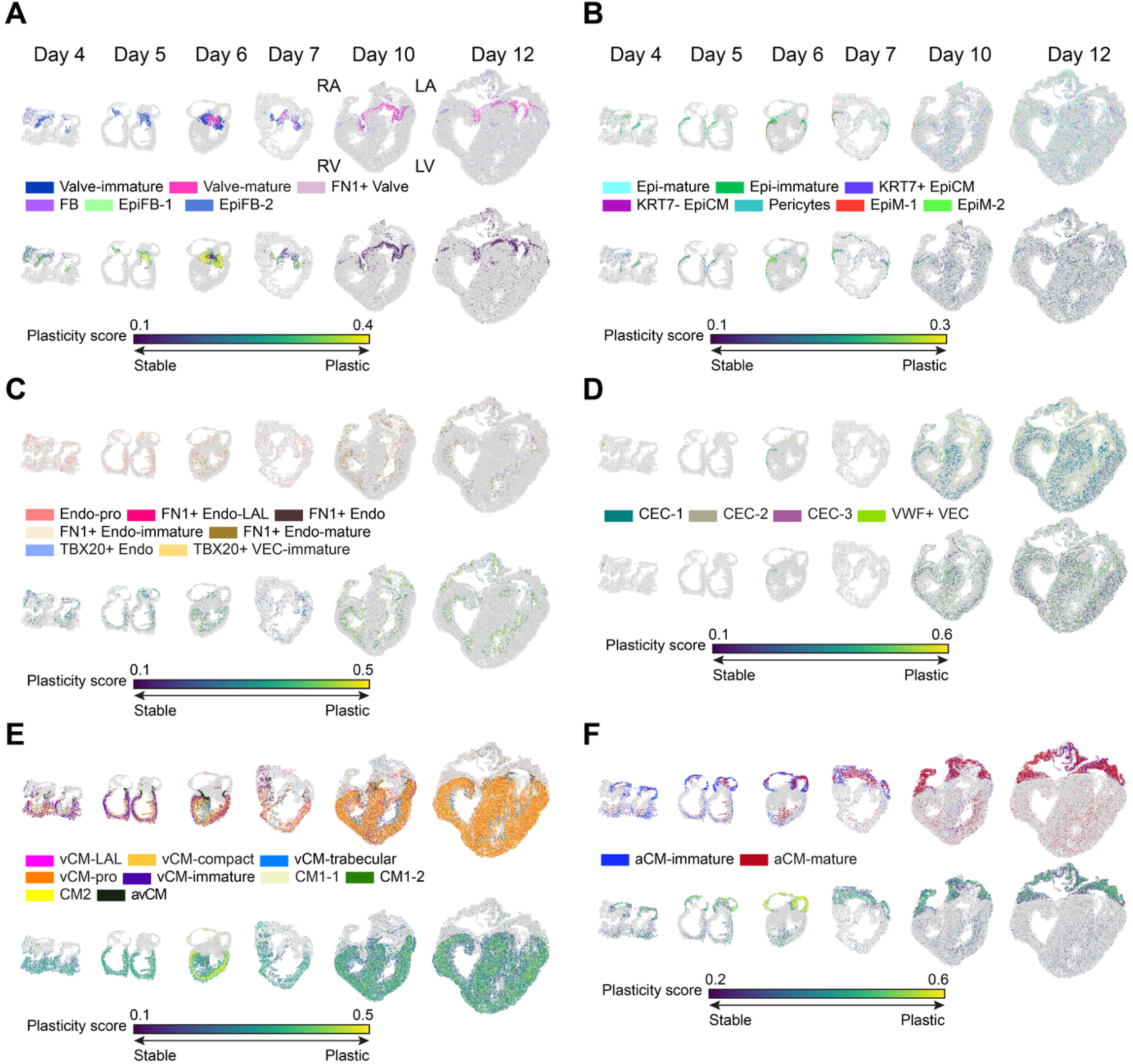
Spatial Cell type map and Plasticity score map in Normal heart. (A-F) Spatial plots of cell types (top) and plasticity scores (bottom) across developmental time points (Day 4, 5, 6, 7, 10, and 12) in normal hearts for (A) fibroblasts, (B) epithelial cells, (C) endocardial cells and TBX20+ VEC, (D) coronary endothelial cells (CEC) and VWF+ VEC, (E) cardiomyocytes excluding aCMs, and (F) aCMs.

**Figure S3.**
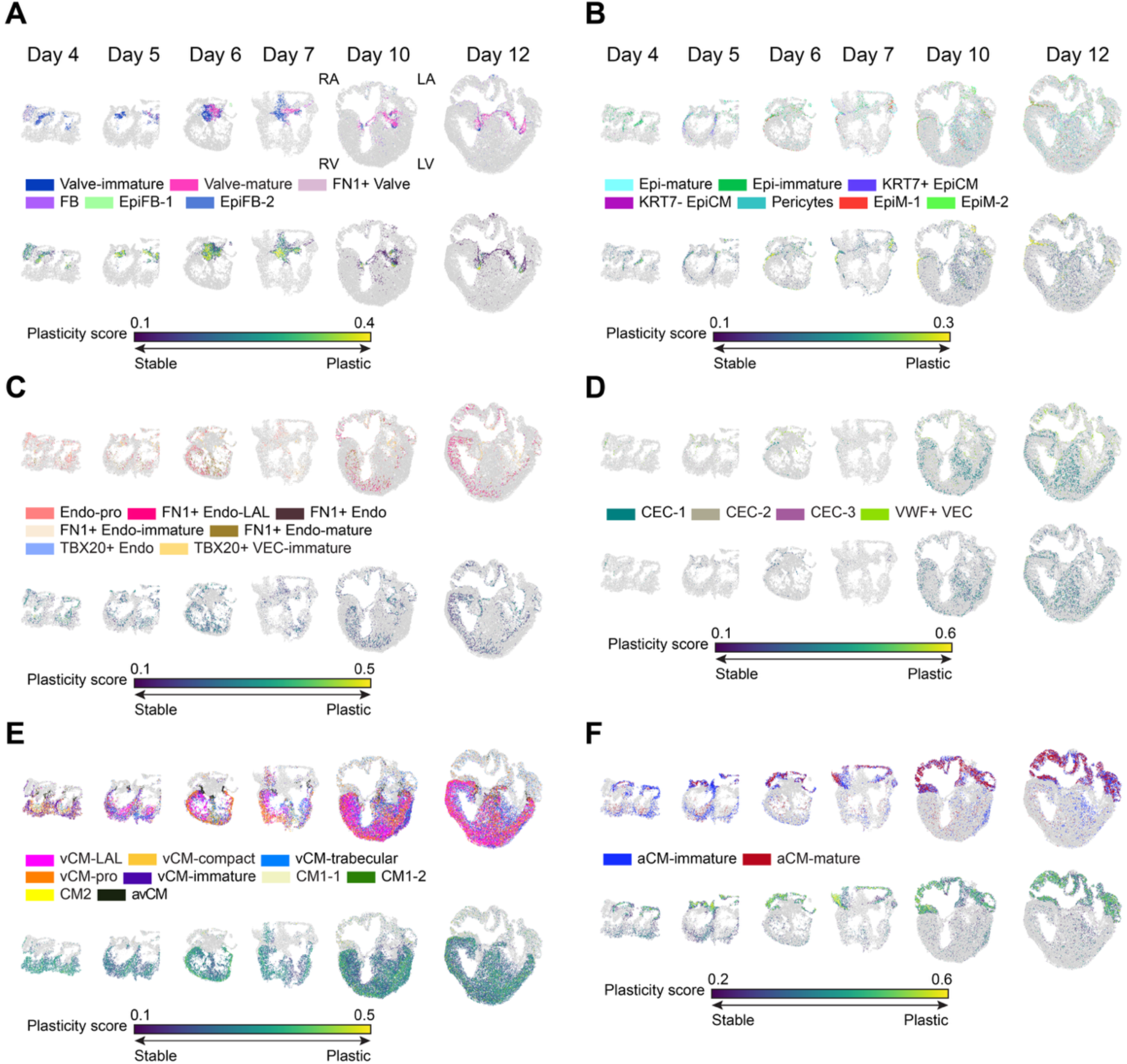
Spatial Cell type map and Plasticity score map in LAL heart. (A-F) Spatial plots of cell types (top) and plasticity scores (bottom) across developmental time points (Day 4, 5, 6, 7, 10, and 12) in LAL hearts for (A) fibroblasts, (B) epithelial cells, (C) endocardial cells and TBX20+ VEC, (D) coronary endothelial cells (CEC) and VWF+ VEC, (E) cardiomyocytes excluding aCMs, and (F) aCMs.

**Figure S4.**
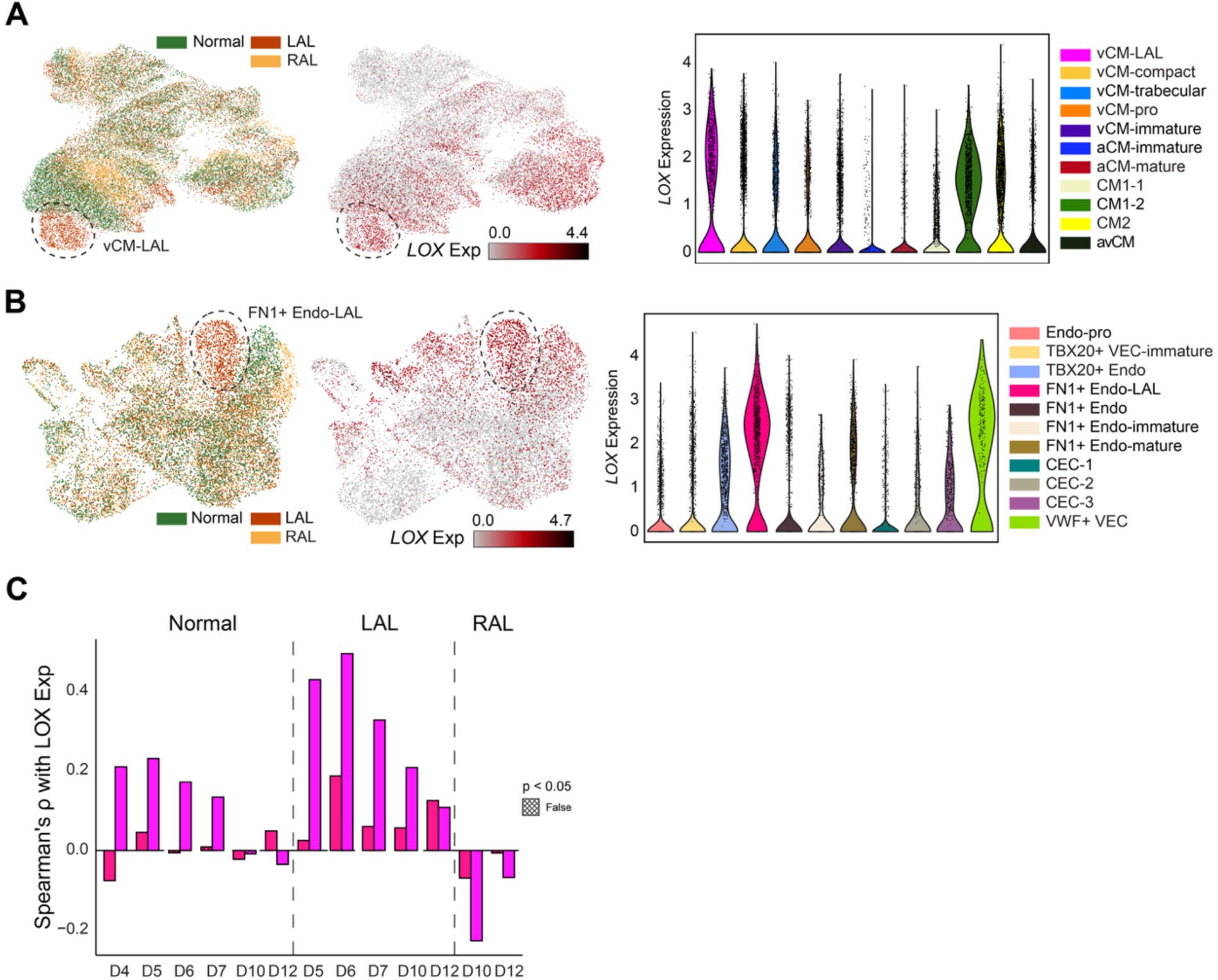
LOX Expressing LAL-enriched cell type. (A) UMAP plot of cardiomyocytes from scRNA-seq data, colored by condition and *LOX* expression. The vCM-LAL cluster is outlined with a dashed line (left). Violin plot of LOX expression (right). (B) UMAP plot of endothelial cells from scRNA-seq data, colored by condition and *LOX* expression. The FN1+ Endo-LAL cluster is outlined with a dashed line (left). Violin plot of LOX expression (right). (C) Bar plot of Spearman’s ρ values showing correlations between *LOX* expression and the proportion of vCM-LAL or FN1⁺ Endo-LAL per ST spot.

**Figure S5.**
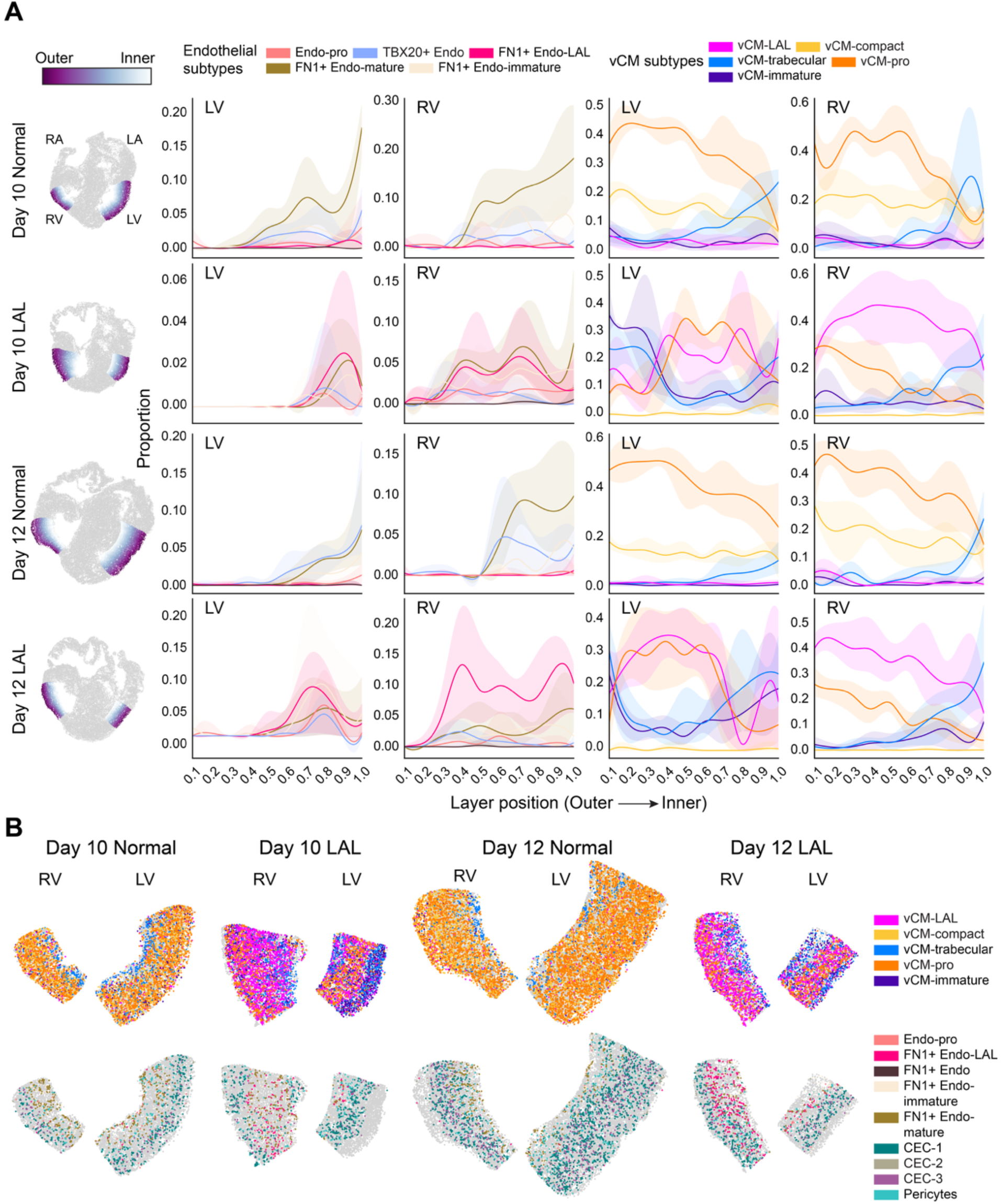
Identification of LOX expressing LAL-enriched cell types and Their Preferential Localization in the RV. (A) Spatial plots of Day 10 and Day 12 normal and LAL hearts colored by layer position (left). Distribution plots showing layer-wise changes in non-vascular endothelial cell and vCM proportions from outer to inner layers of the LV and RV (right). Lines represent mean proportions across five upper-bottom layer bins (**STAR Methods**), and shaded areas indicate 95% confidence intervals. (B) Zoomed in spatial maps of cell types for LV and RV on Day 10 and 12, under different conditions.

**Figure S6.**
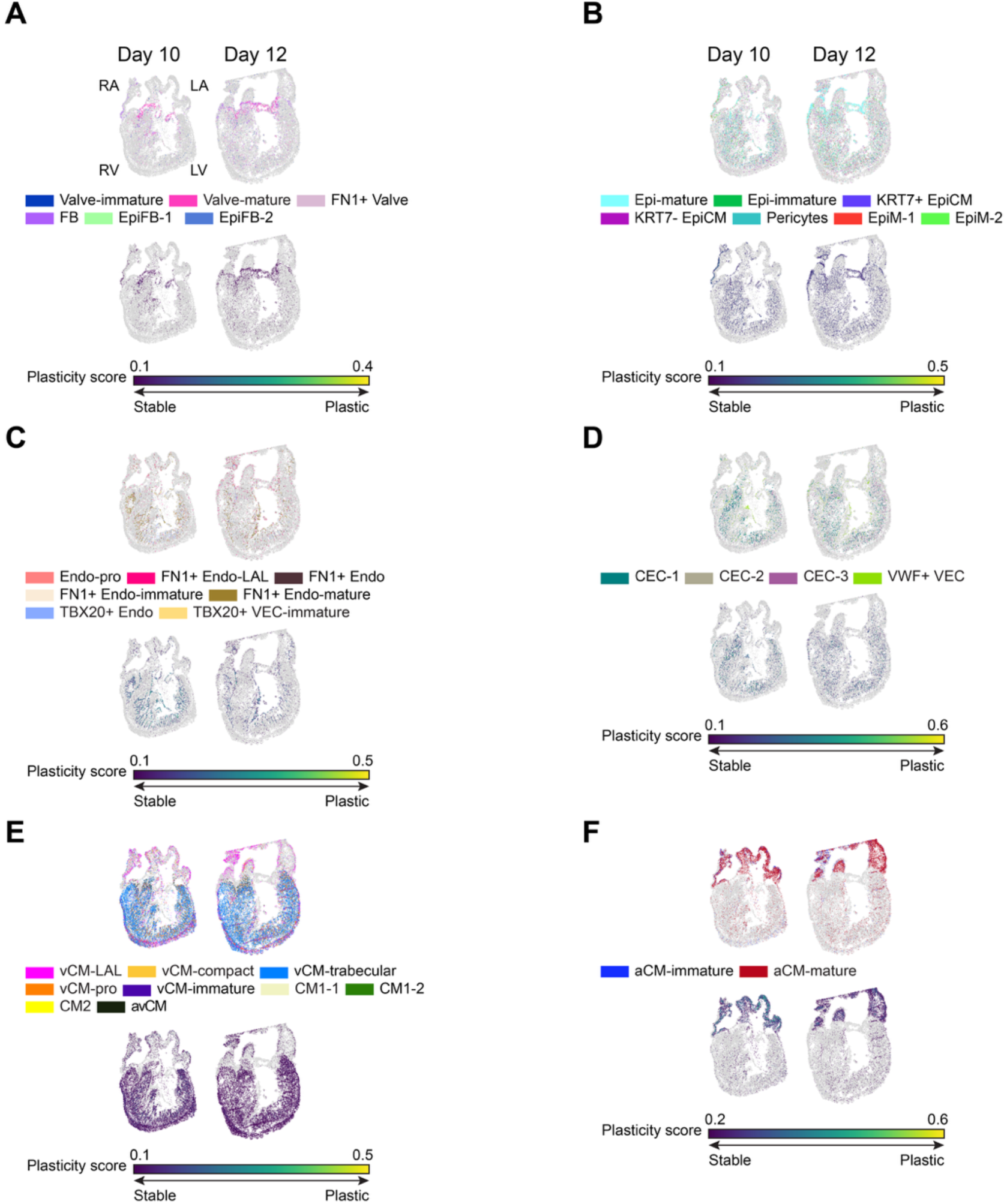
Spatial Cell type map and Plasticity score map in LAL heart. (A-F) Spatial maps of cell types (top) and plasticity scores (bottom) across developmental time points (Day 10 and 12) in RAL hearts for (A) fibroblasts, (B) epithelial cells, (C) endocardial cells and TBX20+ VECs, (D) coronary endothelial cells (CEC) and VWF+ VEC, (E) cardiomyocytes excluding aCMs, and (F) aCMs.

